# Vision reconstructs the cellular composition of binocular circuitry during the critical period

**DOI:** 10.1101/2020.05.31.126474

**Authors:** Liming Tan, Elaine Tring, Dario L. Ringach, S. Lawrence Zipursky, Joshua T. Trachtenberg

## Abstract

High acuity binocularity is established in primary visual cortex during an early postnatal critical period. In contrast to current models for the developmental of binocular neurons, we find that the binocular network present at the onset of the critical period is dismantled and remade. Using longitudinal imaging of receptive field tuning (e.g. orientation selectivity) of thousands of layer 2/3 neurons through development, we show most binocular neurons present at critical-period onset are poorly tuned and rendered monocular. These are replenished by newly formed binocular neurons that are established by a vision-dependent recruitment of well-tuned ipsilateral inputs to contralateral monocular neurons with matched tuning properties. The binocular network in layer 4 is equally unstable but does not improve. Thus, vision instructs a new and more sharply tuned binocular network in layer 2/3 by exchanging one population of neurons for another and not by refining an extant network.

**One Sentence Summary:** Unstable binocular circuitry is transformed by vision into a network of highly tuned complex feature detectors in the cortex.

## Introduction

Cortical circuitry in the mammalian brain is especially sensitive to and instructed by environment-specific stimuli (Espinosa and Stryker, 2012; Hooks and Chen, 2020; White and Fitzpatrick, 2007; WIESEL and HUBEL, 1963). The cortex is composed of many different neuron types and each of these forms a dense complex network of synaptic connections (Gouwens et al., 2019; Tasic et al., 2018). Early steps in the establishment of neural circuitry are governed by genetically hard-wired mechanisms mediated by cell recognition molecules, synaptic adhesion proteins, and sensory-independent neural activity (Ackman et al., 2012; Katz and Shatz, 1996; Ko et al., 2013, 2014; Sanes and Zipursky, 2020; Xu et al., 2011). This circuitry is then improved and solidified by experience, a stage in development referred to as the critical period (Espinosa and Stryker, 2012).

The impact of sensory experience on the development of cortical circuitry is perhaps best characterized in the primary visual cortex. Visual information is processed within the retina and then transmitted by retinal ganglion cells to thalamocortical relay neurons. These neurons, in turn, project to the primary visual cortex forming synapses with layer 4 neurons and to a lesser extent with neurons in layer 2/3 (Hooks and Chen, 2020; Niell, 2015). Within cortex, a complex translaminar and intralaminar web of feedforward and feedback synapses establish a new circuitry of neurons whose receptive field complexity gives rise to feature detection not present at antecedent levels of visual processing. This complexity is informed by the activity pattern of sensory receptors in the epithelium and refined by experience (Ackman et al., 2012; Hoy and Niell, 2015; Paik and Ringach, 2011; Ringach, 2004).

The emergence of binocular neurons in cortex is especially sensitive to visual experience during a critical period of early postnatal development. In visual cortex, responses emerge first to contralateral eye stimulation (Crair et al., 1998; Smith and Trachtenberg, 2007). These responses, with the exception of direction tuning (Li et al., 2008; Smith et al., 2015), are innately established, are present at eye opening (Hoy and Niell, 2015; HUBEL and WIESEL, 1963; Ko et al., 2013), and continue to develop along their normal trajectory even in the absence of vision (Ko et al., 2014; Sherk and Stryker, 1976; White et al., 2001). Cortical responses to ipsilateral eye stimulation develop later and their development is regulated by vision (Crair et al., 1998; Faguet et al., 2009; Smith and Trachtenberg, 2007). The most salient feature of critical period plasticity is the experience-dependent establishment of a binocular network composed of neurons whose orientation tuning preferences from each eye are matched. A common view is that intrinsic mechanisms establish a rudimentary binocular circuitry that is subsequently improved by visual experience (Wang et al., 2010, 2013). This is thought to involve gradual refinements in the orientation preferences of intrinsically established binocular neurons, enabling them to respond to the same features viewed by the two eyes (Espinosa and Stryker, 2012).

We longitudinally tracked the tuning properties of binocular and monocular neurons in layer 4 and 2/3 across the classically defined critical period in mouse visual cortex, which spans the fourth and fifth postnatal weeks. These measures show that responses of layer 2/3 binocular neurons become significantly more selective to orientation, develop preferences to higher spatial frequency stimuli, and become more “complex”, or phase invariant. These changes do not occur by gradually improving and matching tuning properties of existing binocular neurons. Instead, the early binocular network in layer 2/3 is remade. More than half of the early binocular neurons are poorly tuned and rendered monocular. In parallel, vision instructs the recruitment of new binocular neurons largely by conversion of the most selective neurons drawn from the monocular contralateral pool. These contralateral neurons gain ipsilateral responses with matched tuning properties. We demonstrate that this transformation in the binocular network requires vision-dependent maturation of cortical responses to the ipsilateral eye, is not driven by changes in layer 4 and is unique to the critical period. Together, these findings define mechanisms by which vision drives the maturation of binocular vision in the primary visual cortex.

## RESULTS

### Receptive field tuning measured from GCaMP6s responses

To obtain a clearer view of the development of binocularity in primary visual cortex, we measured receptive field tuning of pyramidal neurons expressing the genetically encoded calcium indicator GCaMP6s (Chen et al., 2013) in critical period (P22-P36) and adult (>P56) mice (Gordon and Stryker, 1996). For each mouse used in this study, the binocular zone was identified as the overlap of contralateral and ipsilateral eye visual field sign maps obtained from epifluorescence retinotopic mapping of GCaMP6s responses (Figure 1A; Garrett et al., 2014; Kalatsky and Stryker, 2003; Wekselblatt et al., 2016). This map was used to target subsequent high magnification, cellular resolution, resonant scanning 2-photon imaging of GCaMP6s signals (Figure 1B, C). During 2-photon imaging, mice were head-fixed but otherwise alert and free to walk on a circular treadmill. Receptive field tuning for each imaged neuron was estimated from the linear regression of the temporally deconvolved calcium response (Figure 1D) to a sequence of flashed, high-contrast sinusoidal gratings of 18 orientations and 12 spatial frequencies presented at 8 spatial phases for each combination of orientation and spatial frequency (Mineault et al., 2016; Methods). For each imaged cell, a “kernel” plotting response strength across orientations and spatial frequencies is obtained. An example of one such kernel and its emergence as a function of time after stimulus onset is given in Figure 1E (see example of a visually unresponsive neuron in Figure S1A,B). Receptive field tuning kernels whose signals were significantly higher than noise were recovered from approximately 70% of imaged neurons (Figure 1F; see also Methods and Figure S2 for methodological details). This value is in close agreement with electrophysiological measures made from head-restrained, alert mice where 60%-80% of neurons are found to be visually responsive (Hoy and Niell, 2015).

**Figure 1.**
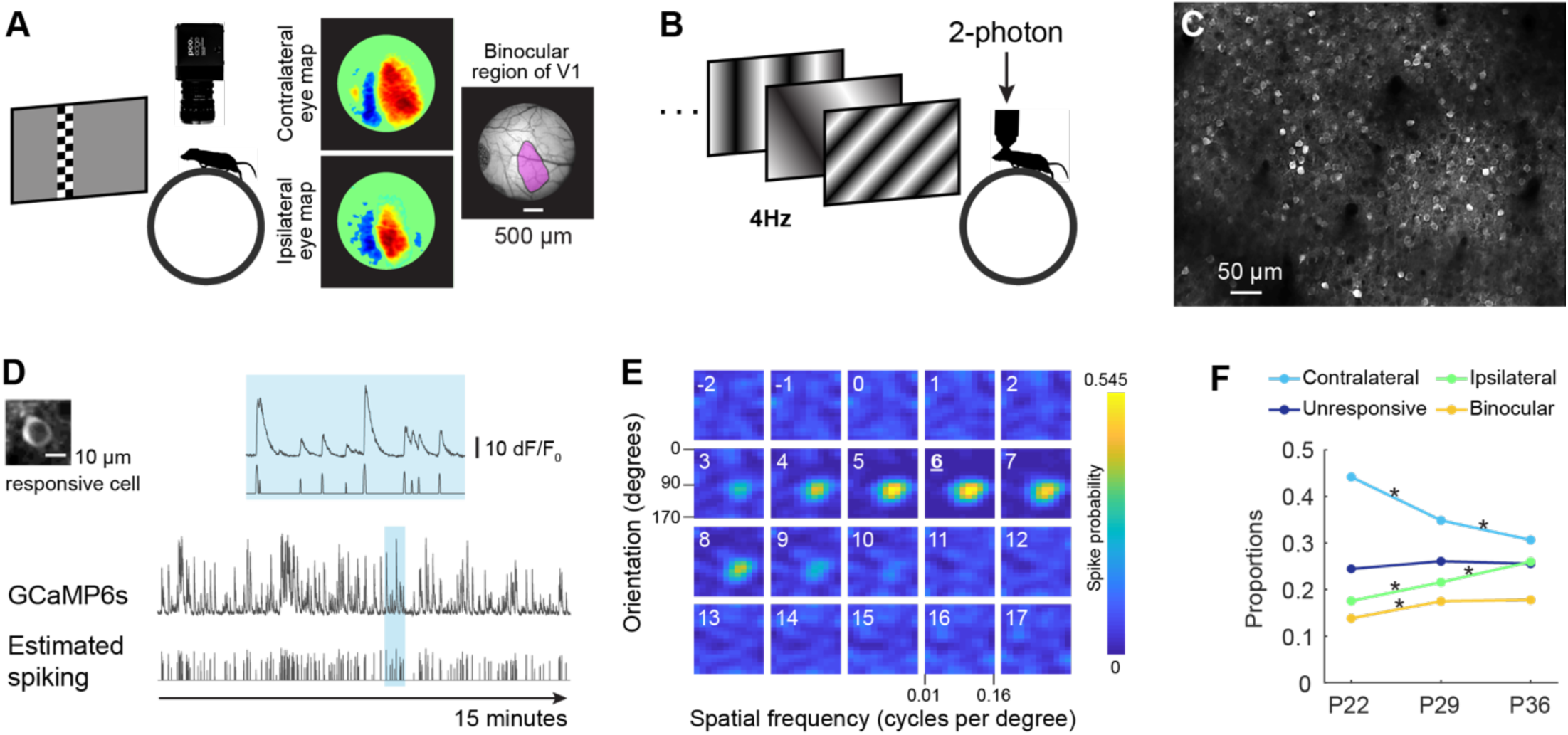
Binocular field mapping and receptive field tuning. **(A)** Visual field maps obtained from low-magnification epifluorescence imaging of GCaMP6s evoked responses to checkerboard bars, which were both drifting and flashing, presented to each eye separately. Red and blue colors in eye-specific maps represent opposite visual field signs designating separate visual areas, with the red sign designating V1 here. The purple shading in the vascular image on the right identifies the binocular region of V1. **(B)** Schematic of 2-photon imaging of visually evoked GCaMP6s responses to a series of sinusoidal gratings sequentially presented at 4Hz. **(C)** Example of a field of view of GCaMP6s labeled neurons in binocular visual cortex. **(D)** Example of fluorescence changes and estimated spike rate over 15 minutes. The cell is shown in the upper left. The narrow blue highlighted region in the full traces has been expanded horizontally and displayed above for clarity. Scale bar for fluorescence changes is shown to the right of this expanded trace. Example is from responses to contralateral eye stimulation. **(E)** Spike probability as a function of stimulus onset for the cell in panel D. Maps are derived from 18 orientations at 12 spatial frequencies presented 16 to 17 times. Numbers in the top left indicate frames relative to stimulus onset. For this cell, peak response occurs 6 imaging frames, or 387 ms after onset of its optimal stimuli. **(F)** Proportions of monocular, binocular and unresponsive neurons as a function of age, for all L2/3 neurons imaged in NR mice across the critical period. * indicate significant changes. Statistics: Tukey’s HSD multiple comparisons test among proportions.

For each neuron, three measures of receptive field tuning were made: 1) orientation selectivity, 2) spatial frequency preference, and 3) complexity. Orientation selectivity was measured as circular variance (see Methods and Figure S1C). Cells that are highly selective, and thus respond to a very narrow range of orientations have lower circular variance. Spatial frequency preference is a measure of spiking as a function of the spacing between lines of the grating. Complexity was measured as the F1/F0 ratio (Skottun et al., 1991; see Methods). So-called “simple” cells respond optimally when a stimulus of a particular orientation and spatial frequency falls within a specific region of that cell’s receptive field. Complex cells, by contrast, are phase-invariant; their responses are unaffected by positional variations of the stimulus within the receptive field (Hubel and Wiesel, 1962, 1968; Ohzawa et al., 1997a; Skottun et al., 1991). Lower F1/F0 values indicate greater complexity and, thus, greater phase invariance. For binocular neurons, a fourth measure was taken: binocular matching. This is a measure of similarity between tuning responses measured through each eye separately. This was calculated as the correlation coefficient of the tuning kernels obtained from each eye (Jimenez et al., 2018). Neurons with similar tuning have higher matching coefficients.

### Selective recruitment and elimination of neurons sharpens the binocular ensemble

To directly examine the development of the binocular network across the critical period in layer 2/3 pyramidal neurons, we made repeated measures of receptive field tuning from the same cells at P22, P29, and P36 (1,064 neurons in four mice tracked across these three time points; Figure 2A; see also Methods and Figure S3 and S4). These measures revealed that the cellular composition of the binocular network changed across the critical period; cells that were binocular at one time point were often monocular at subsequent time points and vice-versa. Of 160 neurons that were binocular at P22, only 64 (40%) remained binocular at P29. This loss was offset by the gain of new binocular neurons; 128 previously monocular or non-responsive neurons became binocular, resulting in a 20% increase in the size of the binocular network from 160 to 192 neurons. This pattern of loss and replenishment repeated in the fifth postnatal week (P29 to 36), though consistent with a tapering of plasticity as the critical period wanes, these dynamics slowed. From P29 to P36, 86 binocular cells were lost, and 91 new binocular neurons were gained. Notably, the number of binocular neurons created by the addition of ipsilateral eye inputs to previously contralateral, monocular neurons was 2.4 times larger than the number created from initially ipsilateral monocular cells gaining contralateral responsiveness (138 vs 57). Examples of ipsilateral and contralateral eye tuning kernels of functionally stable neurons that became binocular and of neurons that became monocular are shown in Figure 2B-D, respectively.

**Figure 2.**
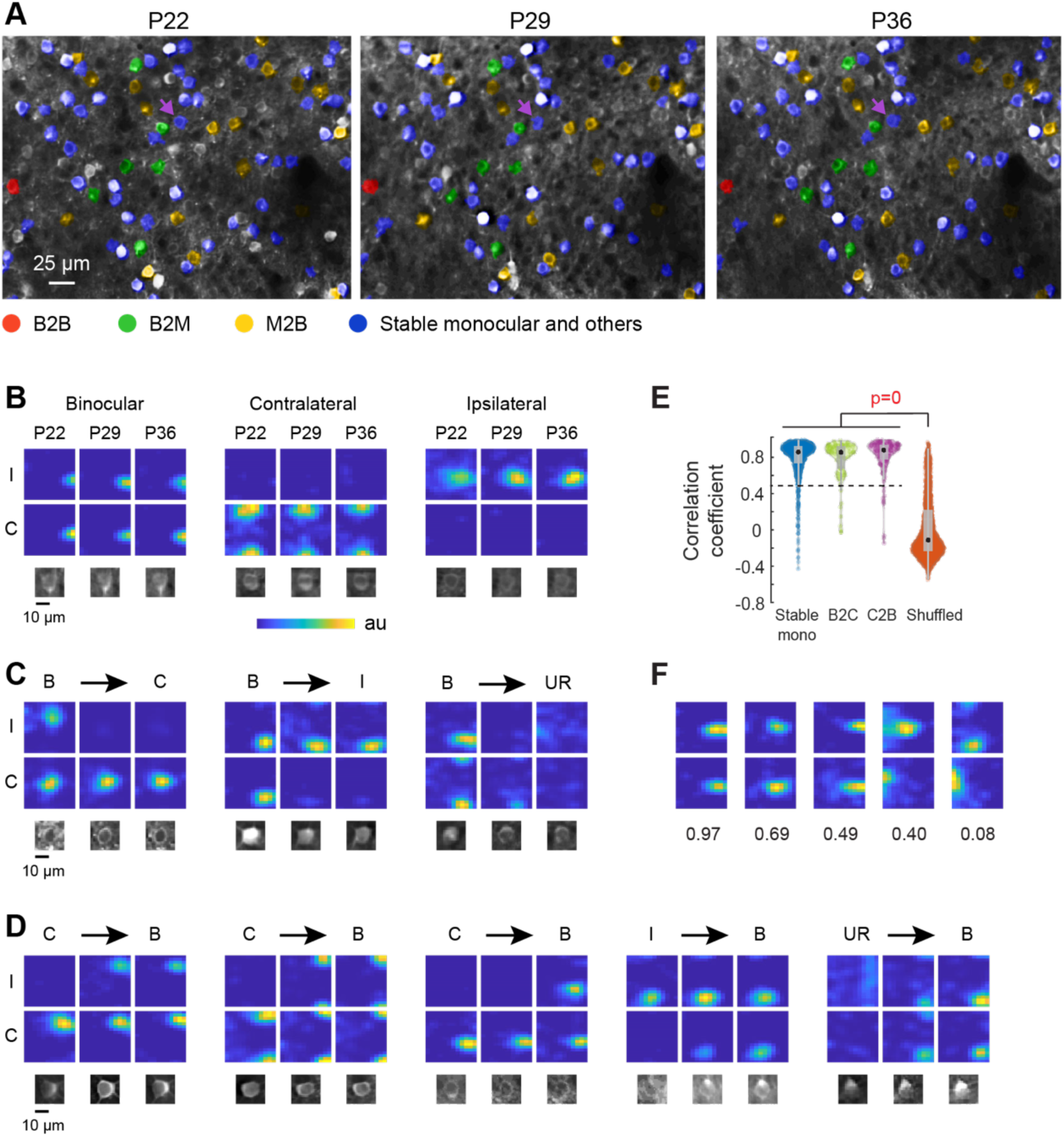
Longitudinal tracking of tuning properties. **(A)** Partial field of view of longitudinal imaging in layer 2/3 across the critical period. Cells tracked for tuning properties across all three time points are colored, with red, green and yellow masks represent stable binocular neurons (B2B), lost binocular neurons (B2M) and binocular neurons recruited from monocular/unresponsive pools (M2B). Blue masks indicate stable monocular neurons or neurons with other trajectories. Cells without color masks (white) are ones with tuning properties not tracked. Purple arrows highlight the stable ipsilateral neuron shown in B. **(B)** Example of tuning kernels of neurons that were stable over all three time points. I, ipsilateral eye stimulation; C, contralateral eye stimulation. Below each kernel is an image of the cell at each time point. Ordinate, orientation from 0° to 170°. Abscissa, spatial frequency from 0.01 to 0.16 cycles/degree. Color indicates estimated spiking at each possible combination of orientation and spatial frequency. Color map of estimated spiking is normalized for each cell. The contralateral neuron has orientation preference close to 0°, thus its kernels are split at the top and bottom. **(C)** Kernels are plotted as in B, but for binocular neurons that become monocular or unresponsive. B, binocular; C, monocular contralateral; I, monocular ipsilateral; UR, unresponsive. **(D)** Kernels are plotted as in B, but for monocular or unresponsive neurons that become binocular. **(E)** Correlation coefficient comparing tuning kernel stability between stable monocular cells (stable mono), binocular neurons that become contralateral (B2C) and contralateral neurons becoming binocular (C2B), and cells that paired by chance across the critical period. The tapered tail extending towards higher coefficients in the shuffled control reflects random pairing of kernels with high circular variance (>0.7) and low spatial frequency preference (<0.02 cycles/degree). Statistical test: Kruskal-Wallis one-way analysis of variance, with significant p-values shown above in red. Black brackets denote significance using multiple comparison test with Bonferroni correction for pairwise comparisons. Stable mono: n = 854; B2C: n = 84; C2B: n = 138; Shuffled: n = 1541. This is similar for dark-reared animals (not shown). **(F)** Matching coefficients for five pairs of tuning kernels, presented in order of decreasing coefficients. The value of matching coefficient is below each pair.

As a further validation of our approach to identifying the same cells at each time point (see also Methods and Figure S5), we examined the similarity of the receptive field tuning kernels of each tracked cell across the three time points. We reasoned that if the same neuron is tracked, its tuning preferences should remain fairly stable. This similarity from one time point to the next can be measured as the correlation coefficient of the tuning kernels obtained at each time point. For monocular neurons, coefficients are obtained by comparing a given neuron’s tuning kernel measured on P22 to that measure on P29, and then again between P29 and P36. For binocular neurons that become monocular, the tuning kernel for one of the eyes can be measured before and after this transition across all 3 time points. The same is true for cells that were monocular and became binocular. In mice, contralateral eye responses are dominant. We, therefore, examined the matching coefficients of contralateral eye responses of tracked cells and compared the distributions of these coefficients to the distribution of coefficients obtained when tuning kernels were randomly matched, as would occur if different cells were inadvertently tracked at each time point. In all cases, the distributions of matching coefficients could not occur by chance (Figure 2E). To illustrate this more clearly, we plot examples of receptive field tuning kernels with coefficients from 0.08 to 0.97 in Figure 2F. Kernels with matching coefficients above 0.5 are qualitatively well matched. Using this as a benchmark, 93% percent (790/854) of stable monocular neurons, 89% (75/84) of binocular neurons that became monocular contralateral, and 93% (128/138) of monocular, contralateral neurons that became binocular had matching coefficients greater than 0.5 (Figure 2E).

These data establish that most binocular neurons present at the beginning of the critical period are lost and replaced by previously monocular neurons that have gained input as the critical period progresses.

### Stable and newly formed binocular neurons are better tuned

We wondered whether receptive field tuning had an influence on which neurons became binocular or monocular. We found that binocular neurons with sharper orientation selectivity, higher spatial frequency preferences, and greater complexity had high binocular matching coefficients. These were significantly more likely to remain binocular than binocular neurons with poorer selectivity, reduced complexity, and poorer matching (Figure 3A-C). Moreover, binocular neurons that lost responsiveness to one eye and were jettisoned to the monocular pool were almost always “simple” cells, while stable binocular neurons were largely “complex” cells (Figure 3D). Simply put, binocular neurons that were poorly tuned and simple tended to have lower binocular matching coefficients. These neurons became monocular. In almost all cases, the strongest input was maintained as the weaker was lost (Figure 3E).

**Figure 3.**
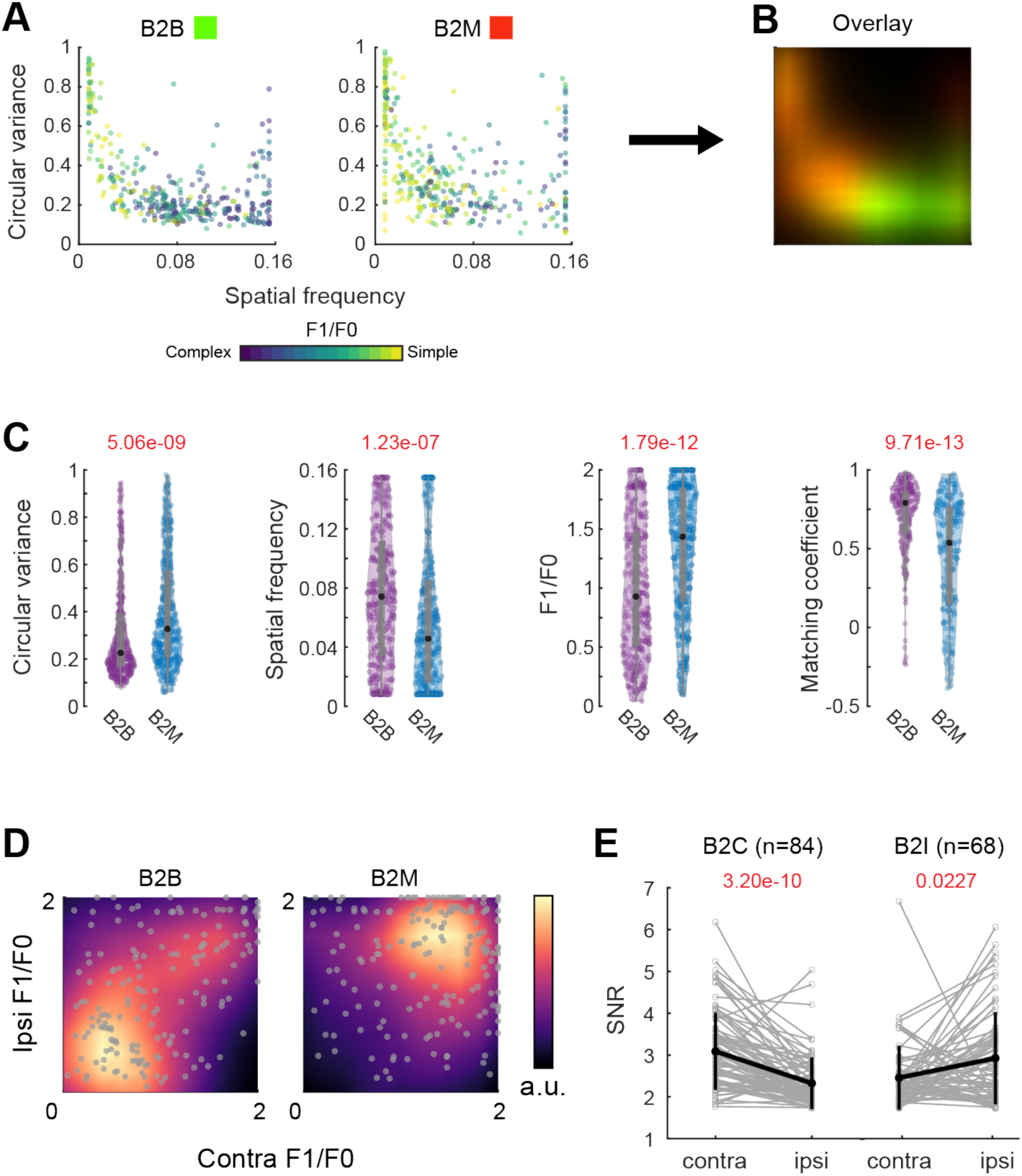
Stable binocular neurons are better tuned during the critical period. **(A)** Tuning measures for binocular neurons retained or lost between P22-29 or P29-36. Each dot represents response with circular variance, spatial frequency and F1/F0 (color bar) to contralateral or ipsilateral eye of a binocular neuron prior to the retention (B2B; 170 neurons) or loss (B2M; 182 neurons) of binocularity. **(B)** Overlay of the tuning density profiles of binocular neurons from (A). Cells that remained binocular (green) have higher spatial frequency and lower circular variance than those rendered monocular (red). **(C)** Receptive field tuning measurements and matching coefficients of stable (B2B; 170 neurons) and lost binocular neurons (B2M; 182 neurons), prior to the transition. Black dot, median; gray vertical line, quartiles with whiskers extending to 2.698σ. P-values, Mann-Whitney U-test. **(D)** F1/F0 plots for the ipsilateral and contralateral eye evoked responses of stable (B2B) and lost binocular neurons (B2M) prior to the transition. Color bar, density. Cells in the lower left and upper right quadrants are “complex” and “simple”, respectively. **(E)** SNR of responses to contralateral and ipsilateral eyes (contra and ipsi, respectively) of binocular neurons rendered monocular contralateral (B2C) or ipsilateral (B2I) prior to transition. Gray, measurements for individual neurons; black, mean ± standard deviation. P-values, Mann-Whitney U-test between contralateral and ipsilateral SNR for each group.

A similar analysis of monocular neurons that became binocular neurons showed that these neurons were better tuned than monocular neurons that remained monocular (Figure 4). Monocular neurons that were recruited into the binocular pool were more selective for orientation, had higher spatial frequency preferences, and were more complex than neurons that remained monocular. Notably, the newly formed binocular neurons had better matched tuning kernels for each eye than the binocular cells that became monocular. We did not see the gradual matching of ipsilateral and contralateral eye receptive field tuning properties proposed by existing models.

**Figure 4.**
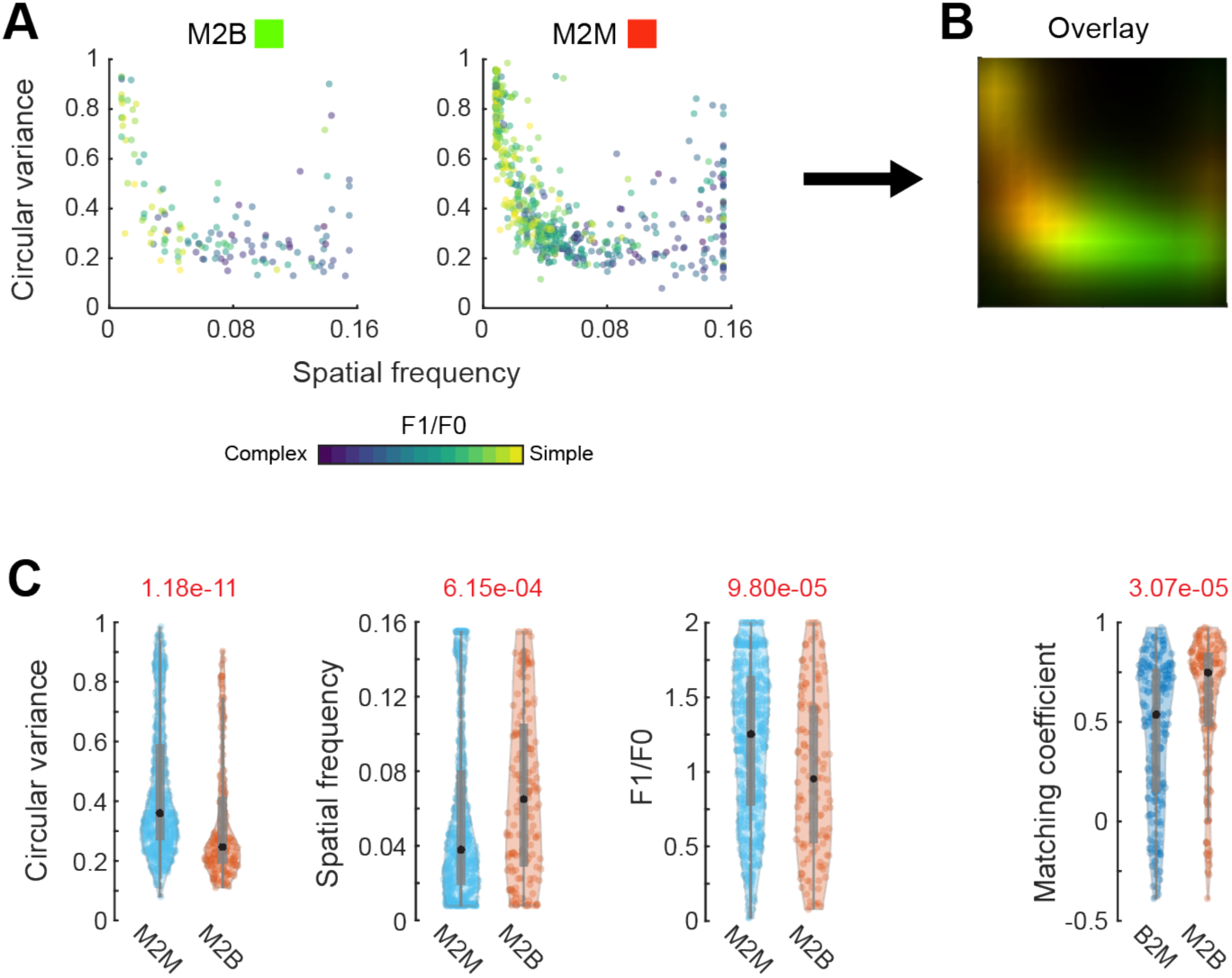
Monocular neurons recruited to binocular pool are better tuned during the critical period. **(A)** Tuning measures for monocular contralateral neurons which became binocular (M2B, 138 neurons) or remained monocular (M2M, 516 neurons) between P22-29 and P29-36. Each dot represents a cell with circular variance, spatial frequency and F1/F0 (color bar), prior to the transition. **(B)** Tuning density profiles of monocular contralateral neurons. Cells that became binocular (M2B, green) have higher spatial frequency and lower circular variance than those that remained monocular (M2M, red). **(C)** Receptive field tuning measurements in panel A, and matching coefficients for newly formed (M2B, n=138) and lost (B2M, n= 182) binocular neurons. Black dot, median; gray vertical lines, quartiles with whiskers extending to 2.698σ. P-values, Mann-Whitney U-test.

The measures indicate that receptive field tuning of the binocular network improves by recruiting the most selective and complex cells from the monocular pool while the broadly tuned and simple binocular cells become monocular. Figure S6 shows tuning measures for each individual mouse. An accounting of the tuning of all cells imaged acutely at P22, P29, and P36 is given in Figure S7 and demonstrates the progressive improvement of binocular and ipsilateral responses across the critical period.

We next set out to ask whether this exchange of neurons and the improvement in their tuning properties required vision, was unique to layer 2/3, and was temporally restricted to the critical period.

### Recruitment but not elimination requires vision

To assess the role of vision in breaking down and re-building the binocular neuron population we measured receptive field tuning and binocular matching in mice that were normally reared until P22 and then placed in the dark from P22 through P36. These mice were imaged prior to dark exposure at P22, again at P29 while they viewed for two hours the high-contrast sinusoidal gratings before returning to darkness, and again at P36. This near complete absence of vision attenuated the normal exchange of neurons between the binocular and monocular networks (three mice, 560 neurons longitudinally tracked). Without vision, receptive field tuning properties of monocular neurons that gained responsiveness to the other eye and became binocular were no better tuned than those that remained monocular, and these newly formed binocular neurons were simpler than monocular neurons that remained monocular (Figure 5A). Thus, vision plays a key role in recruiting complex neurons with improved circular variance and increased spatial frequency into the binocular pool.

**Figure 5.**
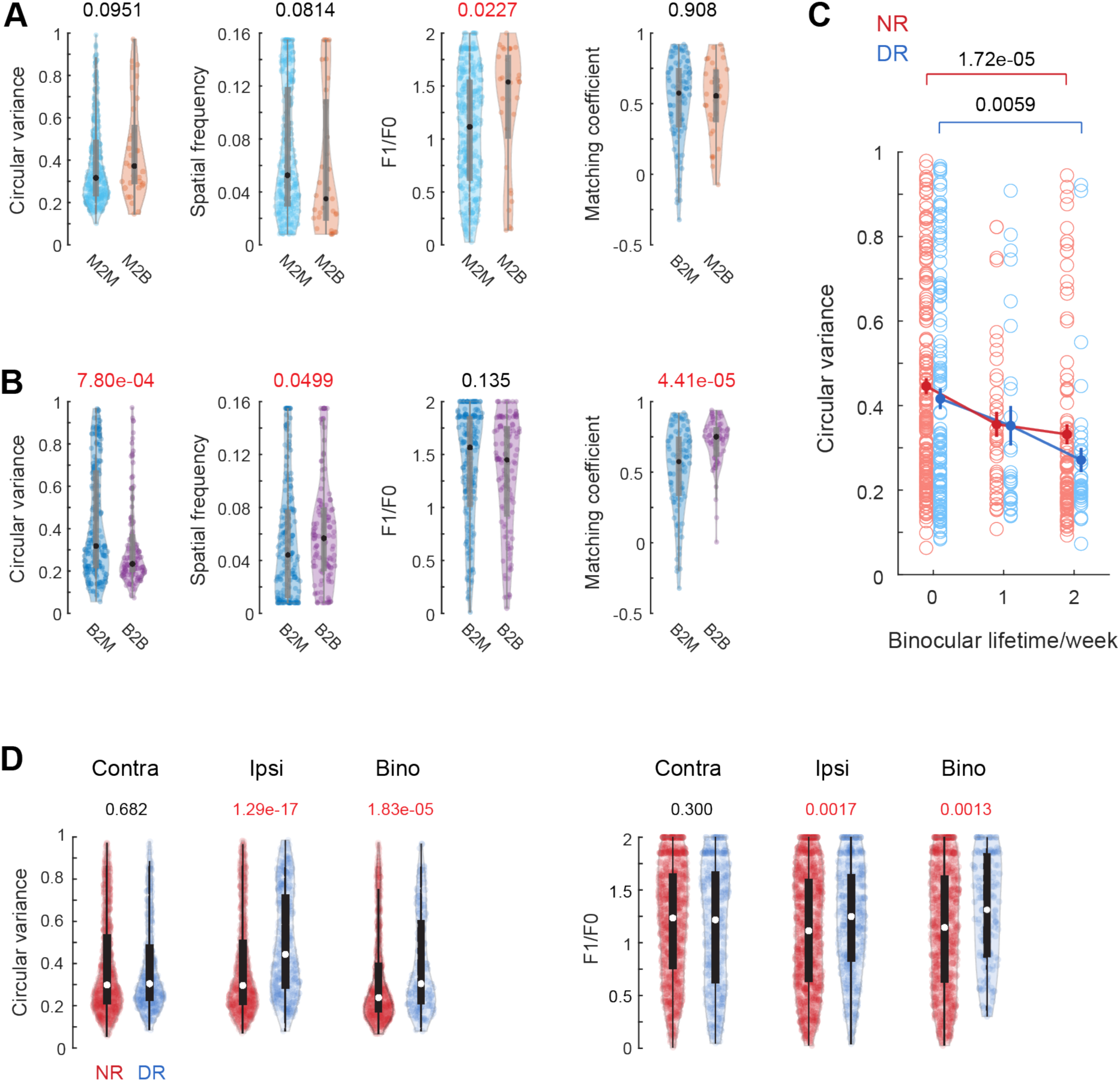
Vision regulates recruitment of better tuned binocular neurons. **(A)** Receptive field tuning and matching coefficient measurements as in Figure 4C for dark reared (DR) mice. M2B, n = 33; B2M, n = 96; and M2M, n = 305. **(B)** Receptive field tuning and matching coefficient measurements as in Figure 3C for DR mice. B2B, n =59; B2M, n = 96. **(C)** Plot of circular variance as a function of binocular lifetime of neurons in normally reared (NR) and DR mice. Each circle represents response to contralateral or ipsilateral eye stimulation of a binocular neuron. Abscissa: 0, neuron was binocular at P22 but not at P29; 1, neuron was binocular at P22 and P29, but not P36; and 3, binocular at all three time points. For each condition (NR and DR), mean+/-SEM is plotted as dot and line at each lifetime. Mann-Whitney U-test with Bonferroni correction (α = 0.0167) was used, with significant p-values shown above their pairwise comparisons. Sample number: NR: 160, 40, 88 from 0 to 2 weeks; DR: 118, 26, 38 from 0 to 2 weeks. No statistical difference exists between the distribution in circular variance for neurons from NR and DR for each binocular lifetime. **(D)** Circular variance and F1/F0 for neurons imaged at P36 in NR and DR mice. “Contra” represents responses to contralateral eye from monocular contralateral and binocular neurons; “Ipsi” represents responses to ipsilateral eye from monocular ipsilateral and binocular neurons; “Bino” represents responses to contralateral or ipsilateral eye from binocular neurons. White dot, median; black vertical lines, quartiles with whiskers extending to 2.698σ. NR Contra, n = 923; DR Contra, n = 404; NR Ipsi, n = 834; DR Ipsi, n = 459; NR Bino, n = 678; DR Bino, n = 156. P-values, Mann-Whitney U-test comparison between NR and DR.

The loss of binocular neurons, however, followed the same pattern as we observed in normally sighted mice. Binocular neurons that became monocular in the absence of vision had broader orientation selectivity, responded to lower spatial frequencies and were more poorly matched than binocular neurons retained (Figure 5B). Indeed, as in normally sighted mice, the stability of a binocular neurons (i.e. their lifetimes) was strongly correlated with circular variance (Figure 5C); neurons with the sharpest tuning were most stable. Figure S8 plots receptive field tuning measures for each mouse separately.

Moreover, ipsilateral eye tuning failed to improve in the absence of vision. Contralateral eye tuning, however, was unaffected (Figure 5D). This immature ipsilateral eye tuning would limit the selectivity and matching of binocular neurons and may explain why the binocular network fails to improve in the absence of vision (see Discussion).

### No improvement in L2/3 in older mice

To determine if the turnover and improvements in the layer 2/3 binocular network are restricted to the critical period, we took repeated weekly measures of receptive field tuning in three mice that were each older than P56 (Figure 6A), which is well past the closure of the critical period in mice. We first looked at the tuning distributions of all cells imaged at each time point. No improvement in the binocular or monocular network was found across either one- or two-week time intervals (Figure S9). Measures of 450 longitudinally tracked cells showed that the degree of turnover was also less in adults (32%, Figure 6B) and, correspondingly, the fraction of stable neurons was higher (62%, 56/91) compared to the critical period (22%, 44/197, p = 8.21e-11, Chi-square test). Moreover, cells that joined the binocular pool were not better tuned than those that became monocular (Figure 6C). These data show that the turnover of binocular neurons is markedly reduced in the adult and, by contrast to the critical period, there was no improvement in the tuning of the binocular network.

**Figure 6.**
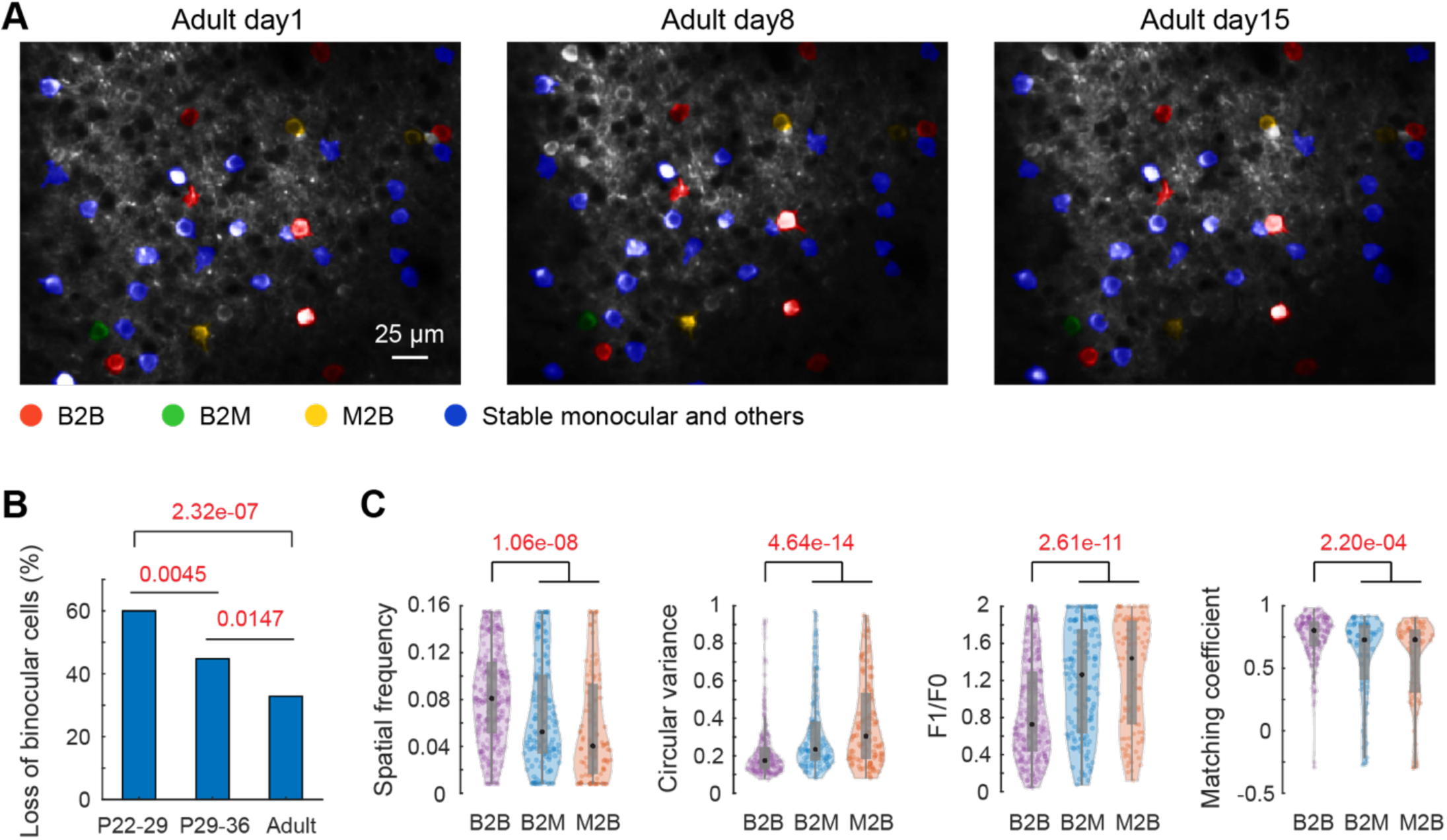
Binocular pools change but tuning properties do not in adult L2/3 neurons. **(A)** Partial field of view of adult layer 2/3 neurons longitudinally imaged. Cells tracked for tuning properties across all three time points are colored, with red, green and yellow masks representing stable binocular neurons (B2B), lost binocular neurons (B2M) and binocular neurons recruited from monocular/unresponsive pools (M2B). Blue masks indicate stable monocular neurons or neurons with other trajectories. Cells without color masks are ones with tuning properties not tracked. **(B)** Increased stability of binocular neurons in adult. The fraction of binocular neurons lost during a week is: B2M/(B2B+B2M)). Chi-square test for pairwise comparisons; P values, red; Bonferroni corrected, α = 0.0167. **(C)** Tuning measurements of contralateral or ipsilateral eye for stable, lost and new binocular neurons (B2B, B2M and M2B, respectively). For B2B and B2M, measurements prior to transition are plotted; for M2B, measurements post-transition are plotted. Black dots, median; gray vertical lines, quartiles with whiskers extending to 2.698σ. B2B: n =137; B2M: n = 67; M2B: n = 54. Statistics: Kruskal-Wallis one-way analysis of variance, with significant p-values shown above each group in red. Black brackets denote significance using multiple comparison test with Bonferroni correction for pairwise comparisons. Note no difference between B2M and M2B.

### No improvement in the layer 4 binocular ensemble

The conventional view of cortical development is that changes in layer 2/3 reflect and amplify changes that occur in the thalamorecipient layer 4 (Freeman and Olson, 1982; Shatz and Stryker, 1978). We therefore checked whether the improvement of the binocular network that we observed in layer 2/3 is also found in layer 4. Measures of receptive field tuning were made at P22, P29, and P36 in mice in which GCaMP6s expression was restricted to layer 4 (717 neurons longitudinally imaged in 4 mice, Figure 7A). The turnover of the binocular network in layer 4 was similar to that in layer 2/3 across the critical period. Of the 717 neurons tracked across the critical period, 155 were binocular at P22. Fifty-seven of these (37%) remained binocular at P29 and 75 new binocular neurons were gained. From P29 to P36 86 binocular neurons were lost as 39 were gained. By contrast, to layer 2/3, however, neither receptive field tuning, nor complexity, nor binocular matching improved (Figure 7B,C; Figure S10). Moreover, all of these measures of receptive field tuning from layer 4 were worse than measures from layer 2/3. Lastly, and again in contrast to layer 2/3, we found no improvement in ipsilateral eye tuning in layer 4 across the critical period (Figure 7D). These data indicate that the improvements in the layer 2/3 binocular network are not inherited from layer 4.

**Figure 7.**
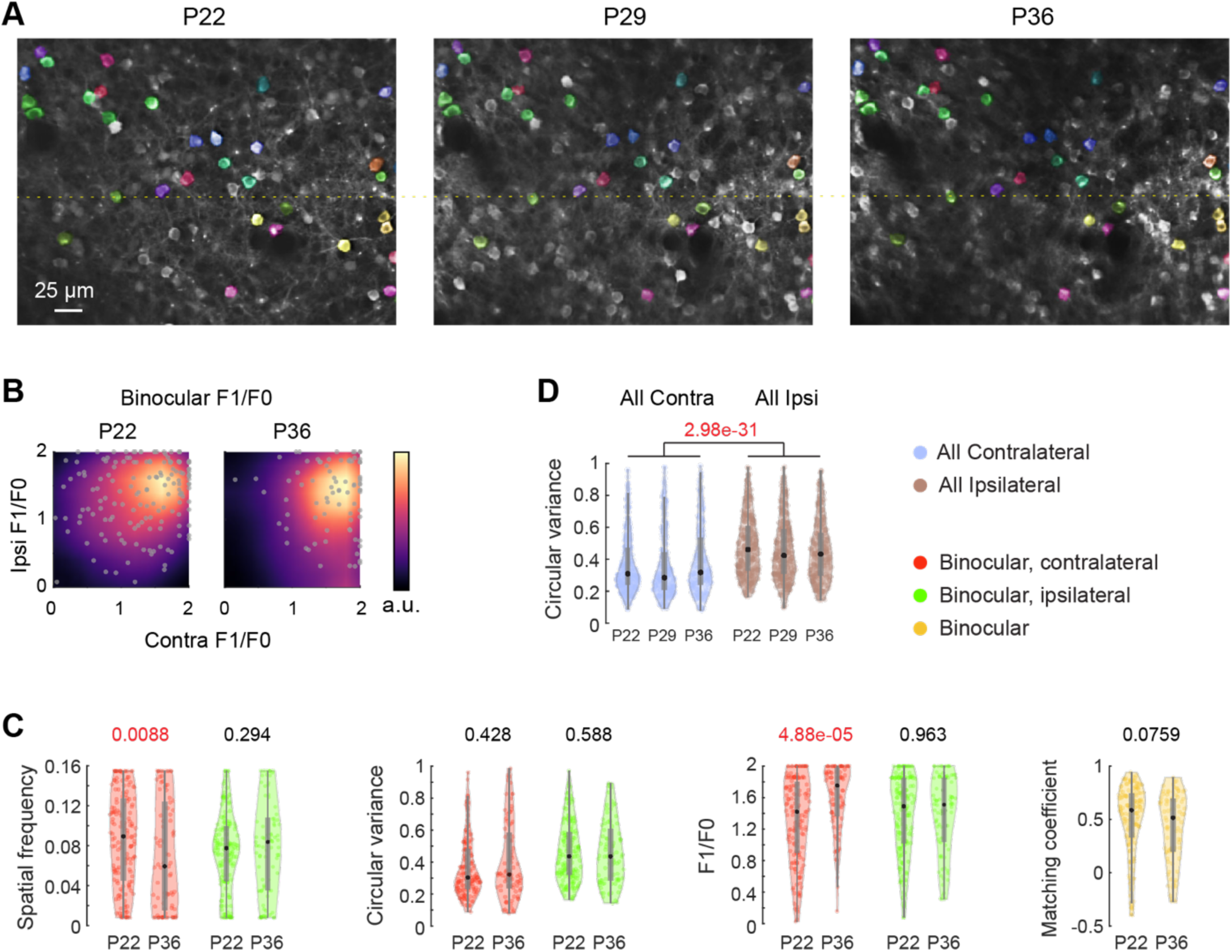
Instability without improvement of layer 4 binocular tuning. **(A)** Partial field of view in layer 4 imaged longitudinally at P22, P29, and P36. Longitudinally tracked cells are colored arbitrarily. A dotted line was added to facilitate visual inspection. **(B)** F1/F0 plots of binocular neurons among longitudinally imaged layer 4 cells at P22 (n=155) and P36 (n=85). Color bar, density (arbitrary units). Upper right and lower left quadrants represent “simple” and “complex” cells, respectively. **(C)** Tuning measurements of binocular neurons at P22 and P36. Color codes are shown to the right of panel D. Black dots, median; gray vertical lines, quartiles with whiskers extending to 2.698σ. P-values (red), Mann-Whitney U-test are shown above each pair of comparisons. **(D)** Circular variance for all contralateral and ipsilateral responses at each age of longitudinally imaged L4 neurons. Statistics: Kruskal-Wallis one-way analysis of variance, with significant p-value shown in red. Black bracket denotes significance using multiple comparison test with Bonferroni correction for pairwise comparisons. P22: 367 contra, 350 ipsi; P29: 278 contra, 352 ipsi; P36, 215 contra, 290 ipsi.

## Discussion

Binocular vision begins to emerge shortly after birth and is actively refined during a critical period of visual cortical development. In humans, this begins about 2-4 months after birth and continues through the first years of infancy and early childhood (Fawcett et al., 2005). In mice, this begins approximately 3 weeks after birth and is complete by the sixth postnatal week (Gordon and Stryker, 1996). Binocular vision and high acuity stereopsis is grounded in the establishment of binocular neurons in primary visual cortex (Ohzawa et al., 1997b). In contrast to the prevailing model (Wang et al., 2010; Xu et al., 2020), the work we present here indicates that binocular neurons present at the onset of the critical period do not undergo improvements in receptive field tuning across the critical period. Instead, longitudinal imaging of identified neurons reveals that the binocular network is particularly unstable during the critical period and that this instability is exploited by vision to build a new and improved binocular network.

Three processes re-make the binocular cell population during this developmental period. First, the majority of binocular neurons prior to the critical period (>60%) are rendered monocular. These binocular neurons are poorly tuned, and vision does not improve their tuning. Second, binocular neurons that are well-tuned at the onset of the critical period are retained. Retention is more efficient with vision. And finally, many well-tuned contralateral monocular neurons acquire matched ipsilateral inputs developed during the critical period to become binocular and this conversion relies on vision. Thus, as a consequence of improved ipsilateral eye tuning, vision instructs the formation of new binocular neurons that reflect this sharper tuning while the initial, more broadly tuned binocular network is largely disassembled.

The instability of the binocular circuitry is not restricted to layer 2/3. Longitudinal imaging of layer 4 cortical neurons during the critical period revealed that these cells are also highly unstable with gain and loss of binocular neurons. By contrast to layer 2/3, however, tuning properties of layer 4 binocular neurons did not improve across the critical period. Thus, while network instability may be necessary to promote improvements in the tuning properties, it is not, in and of itself, sufficient. Furthermore, the improvements in layer 2/3 appear, at least in part, to be independent of changes in layer 4 circuitry (Liu et al., 2008; Trachtenberg et al., 2000).

No improvements in binocular tuning were seen in the layer 2/3 dark-exposure or layer 4 data sets. In both cases, cortical responses to ipsilateral eye stimulation also did not improve. The tuning of binocular neurons is limited by the tuning of its worst inputs. Thus, a parsimonious model for the emergence of sharper binocular tuning in layer 2/3 is that the improvement in ipsilateral eye responses during the critical period drives the improvement of binocular tuning. Monocular neurons that gain input from the other eye will only maintain this new input if it matches the host; postsynaptic responses driven by each will be coincident and thus both will be strengthened in models of timing-dependent synaptic plasticity. Otherwise, the weaker eye’s input is lost. In this manner, a sharper and better matched binocular network emerges as vision improves ipsilateral eye tuning.

Somatostatin-expressing inhibitory neurons (SST cells) in layer 2/3 are well positioned to drive this sharpening. These neurons are centrally involved in synaptic plasticity as they control dendritic spiking and synaptic integration as well as top-down modulation (Urban-Ciecko and Barth, 2016; van Versendaal and Levelt, 2016; Yavorska and Wehr, 2016). Their responses mature during the critical period (Lazarus et al., 2011) and the inhibition provided by SST cells sharpens orientation tuning selectivity and feature coding of pyramidal neurons (Adesnik et al., 2012; Wilson et al., 2012). Moreover, suppressing SST cell activity during the critical period impairs the maturation of ipsilateral eye responses and the emergence of binocularity in layer 2/3 (Yaeger et al., 2019). This suggests a broader view of inhibition during the critical period with fast spiking basket cells inhibiting somatic spiking (van Versendaal and Levelt, 2016) and SST neurons regulating dendritic responses acting together to promote the emergence of a stable binocular network.

In conclusion, the studies we report here indicate that vision, and perhaps sensory experience more generally in other regions of the cortex, reconstructs circuitry during the critical period rather than refines it and does so by exploiting a heightened network instability unique to this period of development. Defining the sources of network instability, the targets of experience at the network, cellular, and molecular levels and the means by which experience integrates one set of inputs into an otherwise hard-wired circuitry are crucial questions for future studies.

## Acknowledgments

We thank all members in the Zipursky lab and Trachtenberg lab. S.L.Z. thanks Tom Mrsic-Flogel and Sonja Hofer for hosting him during a sabbatical during which part of this paper was written.

## Author contributions

Design: L.T., D.L.R., S.L.Z. and J.T.T.; mice breeding and surgery: L.T. and E.T.; experiments: L.T.; analysis and writing: L.T., D.L.R., S.L.Z. and J.T.T.

## Data and materials availability

The databases and corresponding analysis and plotting codes are available upon request.

## Materials and Methods

### Contact for reagent and resource sharing

Further information and requests for resources and reagents should be directed to Joshua Trachtenberg.

### Experimental model and subjects

All procedures were approved by UCLA’s Office of Animal Research Oversight (the Institutional Animal Care and Use Committee, IACUC) and were in accord with guidelines set by the US National Institutes of Health. Normally reared mice were housed in groups of 2-3 per cage in a normal 12/12 light dark cycle. Dark-reared mice were housed in groups of 1-3 per cage in a light-tight cabinet that was additionally shielded with two layers of polyurethane-coated black nylon sheet (Thorlabs, BK5). Animals were naïve subjects with no prior history of participation in research studies. A total of 19 mice, both male (11) and female (8) were used in this study (Critical period layer 2/3 normally reared, 4 males; Critical period layer 2/3 dark reared, 2 males and 1 female; adult layer 2/3, 3 females; critical period layer 4 normally reared, 2 males and 2 females; imaging focal plane displacement, 3 males and 2 females). Mice: All imaging was performed on mice expressing the slow variant of GCaMP6 in pyramidal neurons. For layer 2/3 imaging, these mice were derived from crosses of B6;DBA-Tg(tetO-GCaMP6s)2Niell/J (JAX Stock No: 024742; Wekselblatt et al., 2016) with B6;CBA-Tg(Camk2a-tTA)1Mmay/J (JAX Stock No: 003010; Mayford et al., 1996). For layer 4 imaging, these mice were derived from crosses of B6;C3-Tg(Scnn1a-cre)3Aibs/J (JAX Stock No: 009613) (Madisen et al., 2010) with Ai163 (Daigle et al., 2018) (Gift from Dr. Hongkui Zeng in Allen Institute). Mice expressing both transgenes were identified by PCR, outsourced to Transnetyx (transnetyx.com).

### Surgery

All epifluorescence and two-photon imaging experiments were performed through chronically-implanted cranial windows as in Yaeger et al., 2019. In brief, for critical period experiments, mice at P14-P16 were administered with carprofen analgesia prior to surgery. Mice were anesthetized with isoflurane (5% for induction; 1.5–2% during surgery); body temperature was maintained at 37°C. A thin layer of ophthalmic ointment was applied to both eyes to prevent desiccation. Anesthetized mice were mounted on a stereotaxic surgical stage and the head secured by ear bars and a mouth bar. Scalp overlying the parietal plates on both hemispheres of the skull was removed. After the exposed skull dried, it was covered by a thin layer of Vetbond at the junction of any muscle or skin; this junction was further sealed by a layer of black dental acrylic. A stainless-steel head bar was affixed with dental acrylic caudally to V1. Note the headbar needs to be leveled and parallel to a virtual line connecting the two eyes. A high-speed dental drill was used to remove a 3.5 mm diameter portion of the exposed skull overlying V1 on the left hemisphere; care was taken not to damage the dura. A sterile 3 mm diameter cover glass was place inside the craniotomy to cover the exposed brain and then sealed to the surrounding skull with Vetbond. The edges, as well as the remainder of the exposed skull and surgery margins were sealed with dental acrylic. Mice were recovered on a water-circulating heating pad. When alert, they were placed back in their home cage. Carprofen was administered daily for 3 days post-surgery. Mice were left to recover for 4-7 days prior to imaging. For adult experiments (imaging started at P56, P60 and P77 for each mouse, respectively), surgeries were performed 10-14 days before experiments. For experiments on imaging effects of focal plane displacement (Figure S5), we performed surgery on 5 mice at P18, P19, P27, P37 and P48, respectively.

### Mapping of binocular area of the primary visual cortex

The binocular region of primary visual cortex on the left hemisphere for each mouse used in this study was identified using low magnification, epifluorescence imaging of visually-evoked GCaMP6s signals (Salinas et al., 2017; Wekselblatt et al., 2016). GCaMP6s was excited using a 470nm light-emitting diode. The monitor was positioned relative to each mouse so that the binocular field fell in the middle of the monitor. To map azimuth and elevation we followed a procedure adapted from Kalatsky and Stryker, 2003. We presented a contrast reversing checkerboard (checker size 4×4 deg) windowed by a 1D Gaussian along the horizontal or vertical axis. The Gaussian envelope drifted normal to its orientation to complete a sweep of the entire screen in 10 sec. We used both directions of motion to estimate neural delay and obtain an absolute phase map. The screen size was 120 deg in azimuth and 80 deg in elevation and the monitor was placed 19 cm from the eyes. Eight cycles were recorded for each of the four cardinal directions. Images of GCaMP6s-mediated fluorescent changes were acquired through the camera path of the two-photon microscope (Neurolabware, Los Angeles, CA) equipped with a PCO edge 4.2 m HQ sCMOS camera using a 4X microscope objective (Olympus, 0.16 numerical aperture). Images were acquired at 10 or 15 frames per second. The camera was focused approximately 100 μm below the pia surface. Visual stimulus presentation and image acquisition were controlled by custom written software in Processing sketch using OpenGL shaders (see https://processing.org) and Matlab (Mathworks). Transistor-transistor logic signals generated by the stimulus computer were sampled by the microscope and time-stamped with the frame imaged at that time. This provided the synchronization between visual stimulation and imaging data. Retinotopic maps of azimuth and elevation were generated from these images as in Kalatsky and Stryker, 2003, and visual field sign maps were calculated as in Garrett et al., 2014. The binocular area of the primary cortex was defined as the region of primary visual cortex driven by both the contralateral and ipsilateral eyes (Figure 1A).

### Two photon calcium imaging

2-photon imaging was targeted to the binocular zone of V1 based on the epifluorescence mapping described above. Imaging was achieved using a resonant/galvo scanning two-photon microscope (Neurolabware, Los Angeles, CA) controlled by Scanbox image acquisition software (Los Angeles, CA). GCaMP6s was excited by a Coherent Discovery TPC laser (Santa Clara, CA) running at 920 nm focused through a 16x water-immersion objective lens (Nikon, 0.8 numerical aperture). The objective was set at an angle of 10 to 15 degrees from the plumb line during imaging to reduce the slope of the imaging planes relative to the pia surface. Image sequences (512×796 pixels, 490×630 μm) were captured at 15.5 Hz at a depth of 90 to 320μm below the pial surface for layer 2/3 imaging, and 350 to 500μm below pia surface for layer 4 imaging. All imaging was performed on alert, head-fixed mice that were free to move on a 3D-printed running wheel. A rotary encoder placed with its post through the wheel axel was used to record running. Eye movements and fluctuations in pupil diameter were recorded using a Dalsa Genie M1280 camera (Teledyne Dalsa) fitted with a 740nm long-pass filter to image infrared laser light scattered through the brain and out the pupil. Both the rotary encoder and eye tracking cameras were triggered at scanning frame rate. To measure response properties of neurons to each eye separately, an opaque patch was placed immediately in front of one eye when recording the responses of neurons to visual stimuli presented to the other eye.

### Visual stimulation during 2-photon imaging

Prior to acquiring data sets, the retinotopic position of the imaging field was mapped using a series of small, flashing checkerboard squares to evoke responses. Based on these measures, the position of the monitor was adjusted so that the receptive fields were centered on the center of the screen. The imaging area on the binocular region spanned ∼20 degrees in azimuth and 16 degrees in elevation. Stimulus contrast was set at 80%.

#### Flashing sinusoidal gratings

Visual stimulation for measuring response properties of neurons were the same as in Jimenez et al., 2018. Briefly, a set of sinusoidal gratings with different orientation, spatial frequency and spatial phase were generated in real-time by a Processing sketch using OpenGL shaders (see https://processing.org). The orientation domain was sampled at equal intervals of 10 degrees from 0 to 170 degrees; the spatial frequency domain was sampled in 12 equal steps on a logarithmic scale from 0.0079 to 0.1549 cycles per degree. For each combination of orientation and spatial frequency, spatial phase was equally sampled at an interval of 45 degrees from 0 to 315 degrees. Static gratings with different combinations of orientation, spatial frequency and spatial phase were presented at a rate of 4 Hz in pseudo-random sequence on a screen refreshed at 60 Hz. Imaging sessions were 15 min long (3600 stimuli in total), so each combination of orientation and spatial frequency appeared 16 or 17 times, and each spatial phase for an orientation/spatial frequency combination appeared twice (responses of neurons as a function of spatial phase is used to calculate F1/F0 values). To synchronize visual stimulation and imaging data, transistor-transistor logic signals generated by the stimulus computer were sampled by the microscope and time-stamped with the frame and line number being scanned at that time.

### Analysis of two-photon imaging data

#### Image processing

The pipeline for image processing is described in Ringach et al., 2016, with modifications for processing imaging planes consisting of cells whose visual responses were recorded to two eyes separately. Briefly, movies from the same plane for each eye were concatenated to a single file. Images in this file were aligned to correct for frameshift caused by motion during imaging (Figure S3A). The mean image of each plane generated after alignment was used to find the same cells for longitudinal imaging. For segmentation of pyramidal neurons, a Matlab graphical user interface tool (Scanbox, Los Angeles, CA) was used to define regions of interest (ROI) corresponding to cell bodies. This segmentation tool was first set to collect 600 to 1200 evenly spaced frames from the motion-corrected movie. This represents ∼2-4% of all imaging frames per experiment. These frames were used to calculate pixel-wise correlations of fluorescence changes over time. The upper limit on the number of frames used to calculate this correlation map was determined by GPU performance (Nvidia Quadro M5000). The temporal correlation of pixels was used to define the boundaries of each neuron (Figure S3B). After segmentation, the fluorescence signal for each cell was extracted by computing the mean of the calcium fluorescence within each ROI and subtracting the median fluorescence from the nearby neuropil (Figure S3C,D). Neural spiking was then estimated via non-negative temporal deconvolution of the extracted signal using Vanilla algorithm (Berens et al., 2018). Non-negative deconvolution outperforms supervised algorithms (Pachitariu et al., 2018). After signal extraction and deconvolution, fluorescent signals and estimated spikes for each cell were split into separate files, corresponding to the separate imaging sessions for the two eyes. In this way, a single index was assigned to each cell, whose responses to each eye were calculated separately, to measure receptive field tuning properties and to define cells as binocular, contralateral monocular, ipsilateral monocular, or visually unresponsive. Each imaging experiment was independently segmented. Segmentation also gave rise to a mask file (796×512 matrix) in which ROIs were labeled by their index numbers; these were used to track the same cells imaged across days (Figure S3B).

#### Calculation of response properties

##### Signal to noise ratio (SNR) of visual response and identification of visually responsive neurons

SNR was used to identify neurons with significant visual responses. First, reverse correlation (Jimenez et al., 2018; Ringach et al., 1997, 2016) of spiking estimation relative to stimulus onset was used to locate the optimal delay for preferred stimuli (Figure 1E). The optimal delay is the frame at which the spiking standard deviation reached its maximum, which indicates the timing of each cell’s peak responses as a function of stimulus onset. This occurred five to seven frames after stimulus onset. SNR for each neuron was calculated based on this timing, with signal being the mean of spiking standard deviation at 5-7 frames after stimulus onset, and noise as this value at frames well before or after stimulus onset (frames –2 to 0, and 13 to 17). Second, this measure of SNR as well as optimal delay were used to score cells as visually responsive or unresponsive. Optimal delay as a function of SNR is plotted for all cells imaged in this study (Figure S2A). Neurons whose optimal delays occurred outside of the time-locked stimulus response window of 4 to 8 frames (padded by ±1 frame around the 5-7 frame range used above; Figure S2A, blue highlight) were spontaneously active, but otherwise visually unresponsive (see Figure S1A,B for example of a visually unresponsive neuron). They had low SNR values (close to 1) and their optimal delays were not time locked to any stimulus. The SNR values of these unresponsive neurons were normally distributed (mean = 1.0) over a narrow range (Figure S2B, blue shaded histogram). Since spontaneously active but otherwise visually unresponsive neurons could have optimal delays at any imaging frame relative to stimulus onset, some will naturally occur in the 4-8 frame time window when visually driven responses also occur (Figure S2B, red shaded). These can be distinguished from visually responsive neurons by SNR. This threshold is defined by the distribution of SNR values obtained above from the spontaneously active neurons. As a conservative measure we set this threshold at 3 standard deviations above the mean SNR of the normal distribution (Figure S2B, vertical black line). Visually responsive neurons had optimal delays between frames 4 and 8, and SNRs greater than this cutoff. The fraction of visually responsive neurons obtained with this approach matches that obtained with electrophysiology measures of neural spiking in alert mice (Hoy and Niell, 2015). SNR values were calculated separately for responses to the ipsilateral or contralateral eye (Figure S1C). These plots for all cells imaged at P22, P29 and P36 are given in Figure S2C-H; note the similarity across ages.

##### F1/F0 measurement for phase-invariance

The modulation ratio, F1/F0, was used to measure phase invariance of each neuron’s response. The sinusoidal visual stimulus gratings used in this study were presented at different spatial phases (Ringach et al., 2002). F1/F0 is the ratio of the amplitude of the post-stimulus time histogram (F1, 1st Fourier harmonic) for a given cell and its mean firing rate (F0, 0th Fourier harmonic). For complex cells the amplitude of F1 is smaller than F0 (F1/F0 < 1), while for simple cells this relationship is inverted (F1/F0 > 1) (Skottun et al., 1991).

##### Tuning kernel for orientation and spatial frequency

The estimation of the tuning kernel was performed as in earlier studies by fitting a linear model between the response and the stimulus (Ringach et al., 2016). Cross-correlation maps were used to assess each neuron’s spiking level to each visual stimulus (orientation and spatial frequency) (Figure 1E). Cross-correlation maps were computed by averaging responses over spatial phases. The final tuning kernel of a neuron was defined as the correlation map at the optimal delay (Figure 1E, frame 6). Orientation and spatial frequency preference, circular variance and binocular matching coefficient were all calculated using tuning kernels for neurons whose SNR exceeded the noise threshold. Example kernels are given in Figure S1C.

##### Orientation and spatial frequency preference

Orientation and spatial frequency preferences were calculated using horizontal (for spatial frequency) and vertical (for orientation) slices of the tuning kernel passed through the peak response. That is, for any given cell, the slice for orientation is a 1×18 array, On, in which a level of estimated spiking (O1 to O18) occurs at orientations θn (0 to 170 degrees, spaced every 10 degrees). Similarly, the slice for spatial frequency is a 1×12 array, Sfk, in which a level of estimated spiking (Sf1 to Sf12) occurs at spatial frequencies ωk (12 equal steps on a logarithmic scale from 0.0079 to 0.1549 cycles per degree). Preferred orientation and spatial frequency were computed as the center of mass of the slices as: 

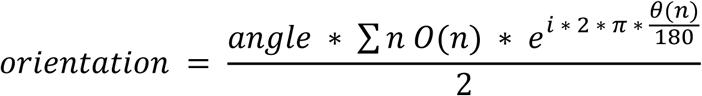

Orientation calculated in this formula is in radian, and was further converted to degrees. 

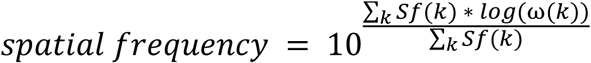

##### Circular variance

The circular variance, *cv*, of a neuron with estimated spiking of On at orientations θn (0° to 170°, spaced every 10°), is defined by 

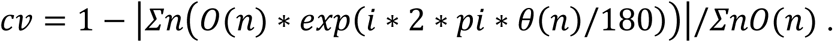

Circular variance is a measure of orientation selectivity, with limits at zero and one. Neurons that do not have orientation tuning have a circular variance of 1. Neurons that are highly orientation selective have circular variance values close to 0 (see examples in Figure S1C).

##### Binocular matching coefficient

The matching coefficient for binocular neurons was defined as the correlation coefficient between their contralateral and ipsilateral tuning kernels.

#### Longitudinal imaging

The same imaging plane was located across days using images acquired on previous imaging days as reference. These include the vascular map at the pia acquired through the PCO camera and the mean motion-corrected fluorescence images of the two photon imaging experiments. Depth of each imaging plane from the pial surface was also written down to facilitate same imaging plane identification. Care was taken to ensure that the mount used to affix each mouse under the microscope objective lens was unchanged in angle or position. During subsequent imaging sessions, using the same objective angle as previous sessions, the approximate region for each imaging plane was first found by matching the imaging region with the corresponding vascular map reference and depth from pia. Fine adjustment of imaging depth was made by manual z-scanning in small steps of 2 μm for ∼20 μm while running visual stimulation to identify the imaging plane that matched the mean image of the same plane acquired previously. For layer 4 imaging, the tdTomato image for each plane was also used to match imaging planes. Imaging planes were considered to be near identical when the position and diameter of the radial vasculature patterns matched, when cell morphology across the four quadrants of the image matched, and when the spatial clustering of local groups of cells across the image matched. An examination of the impact of small variations in focal plane on receptive field tuning and binocularity is given below.

#### Finding longitudinally imaged cells

During segmentation, a single index number was assigned to each segmented neuron.

Since segmentation was performed independently to experiments acquired on different days, the same neurons tracked often had different index numbers across days. Thus, finding the same cells imaged across multiple days involved matching the index numbers for experiments on different days for tracked neurons. This was achieved by assembling an index matrix for cells tracked across multiple days. To identify neurons tracked between adjacent days, a control point-based affine geometric transformation (Matlab syntax: cpselect, fitgeotrans and imwarp) was performed to correct the plane rotation (Figure S4A-C). This transformation was applied to the mask of the cells imaged on the second time point. The overlap of ROIs between the mask file from the first imaging time point and transformed mask file from the second imaging time point was calculated. If the overlapping area between two ROIs A and B was bigger than 50% of the union area of A and B (A∩B > 0.5*(AUB)), the two ROIs were considered to be the same cell (Rose et al., 2016) (Figure S4D-G). The index numbers in each experiment were paired for such cells. Failure to track cells across imaging days could be due to (1) shifts on the x, y axis in the imaging field; (2) cells not segmented in one day but segmented in the other day; (3) cells out of focus in one day and (4) same cells but ROI intersection <50% of ROI union (Figure S4J). Reason (2) accounts for most of the differences between ROI maps, and this is because only 2-4% of the full imaging movie was used to build the pixel-wise correlation maps used for cell segmentation (again, limited by GPU performance). Because the visual stimulus movies are presented in pseudo-random sequence, different sequences will be viewed at one day or the next. Thus, some visually responsive cells may not have responded during the roughly 800 frames used to compute the correlation maps. After the cells tracked in P22-P29 and P29-P36 were identified, cells tracked across P22-P29-P36 were identified by intersecting P22-P29 and P29-P36 index pairs (Figure S4H, I), and index numbers in each day for these cells were concatenated as a matrix. Overall, 56 ± 6% of cells segmented on P36 were also segmented on P22 and P29 in normally reared mice, 47 ± 9% of cells segmented on P36 were also segmented on P22 and P29 in dark-reared mice. This is a significantly better than the 35 ± 3% in Wagner et al., 2019.

#### Analysis of longitudinal data

Using the index matrix, each neuron’s functional class (unresponsive, monocular and binocular) and receptive field tuning properties measured for each eye and across days was assembled as a separate matrix.

#### Experiments on effects of imaging focal plane displacement on the overlap of masks of segmented cells and on receptive field tuning

The effect of focal plane offsets on our measures of receptive field tuning and binocularity is examined in Figure S5. In these control experiments, consistency in segmenting cells and in their receptive field tuning was measured as the focal plane was shifted up or down in 5μm steps over a range of 15μm. This was done across five mice. Consistency in segmenting cells refers to the overlap in segmented cells between two focal planes. For example, segmenting cells at one focal plane and then at a second focal plane above or below the original will give two segmentation masks. The overlap of these two masks will become progressively smaller as the difference in focal plane offset increases. This overlap is measured as the Jaccard index, which is the size of the intersection of these two masks divided by the size of their union. As an example, consider an experiment in which 100 cells are segmented from one focal plane and 110 cells are segmented from another, and 80 of these cells are found to overlap between the two experiments; the Jaccard index is 80/(80+(100-80)+(110-80)) = 0.615.

The Jaccard index and measures of receptive field tuning stability as a function of focal plane offset were made from a total of 9 experiments using 5 mice (details are shown in the table below). In each experiment, measures of visually evoked responses were consecutively taken at 4 focal planes. For four experiments (2, 5, 7, 9 in the table below), response measurements were first taken at focal plane N. These measures were then repeated again at focal plane N (named N’ hereafter) and then at focal planes N–5μm and N+10μm. Thus, six comparisons across focal planes can be considered: one comparison of delta = 0μm (N to N’), two comparisons of delta = 5μm (N to N–5μm and N’ to N–5μm), two comparisons of delta = 10μm (N to N+10μm and N’ to N+10μm), and one comparison of delta = 15μm (N–5μm to N+10μm). To minimize experimental design bias and to collect sufficient data for delta = 0μm and delta = 15μm, we performed additional experiments at N, N’, N+5μm, N–10μm (experiments 3 and 8), N, N’, N– 5μm, N–10μm (experiment 1), and N, N’, N”, N+/–15μm (experiments 4 and 6, respectively). Comparisons of different delta in these experiments are made with the same logic. These comparisons from 5 mice are listed here:

| Experiment | mouse | Imaging age | Depth in 1<sup>st</sup>/<br>μm | Depth in 2<sup>nd</sup>/<br>μm | Depth in 3<sup>rd</sup>/<br>μm | Depth in 4<sup>th</sup>/<br>μm | Data points of pairwise cell intersection/union |
| --- | --- | --- | --- | --- | --- | --- | --- |
| 1 | 1 | P41 | -224 | -224 | -229 | -234 | 1 $\Delta=0$, 3 $\Delta=5$, 2 $\Delta=10$ |
| 2 | 2 | P53 | -204 | -204 | -209 | -194 | 1 $\Delta=0$, 2 $\Delta=5$, 2 $\Delta=10$, 1 $\Delta=15$ |
| 3 | 3, plane1 | P21 | -167 | -167 | -162 | -177 | 1 $\Delta=0$, 2 $\Delta=5$, 2 $\Delta=10$, 1 $\Delta=15$ |
| 4 | 3, plane2 | P22 | -186 | -186 | -186 | -171 | 3 $\Delta=0$, 3 $\Delta=15$ |
| 5 | 4, plane1 | P22 | -179 | -179 | -184 | -169 | 1 $\Delta=0$, 2 $\Delta=5$, 2 $\Delta=10$, 1 $\Delta=15$ |
| 6 | 4, plane2 | P23 | -149 | -149 | -149 | -164 | 3 $\Delta=0$, 3 $\Delta=15$ |
| 7 | 5, plane1 | P31 | -193 | -193 | -198 | -183 | 1 $\Delta=0$, 2 $\Delta=5$, 2 $\Delta=10$, 1 $\Delta=15$ |
| 8 | 5, plane2 | P32 | -222 | -222 | -217 | -232 | 1 $\Delta=0$, 2 $\Delta=5$, 2 $\Delta=10$, 1 $\Delta=15$ |
| 9 | 5, plane3 | P33 | -163 | -163 | -168 | -153 | 1 $\Delta=0$, 2 $\Delta=5$, 2 $\Delta=10$, 1 $\Delta=15$ |
| | | | | | | Total : | 13 $\Delta=0$, 15 $\Delta=5$, 14 $\Delta=10$, 12 $\Delta=15$ |

#### Figure plotting

##### Density profile plots

The code for calculating density profiles was modified from Matlab code scattercloud (https://www.mathworks.com/matlabcentral/fileexchange/6037-scattercloud). Briefly, we first made same number of bins (n=11∼16) along both the x- and y-axis, for pairs of measurements we used as scatter plot. We then calculated density of data points in each bin to get an n-by-n density profile matrix and plot the matrix using Matlab surf function with interpolated coloring for each face. Then we made scatter plot on top of the density profile plot.

##### Density profiles overlay

We overlaid pairs of density profiles by using Matlab imfuse function. Before overlaying two matrices, we normalized each matrix to make the maximum density to be 1 and minimum density to be 0, so that two density profiles being merged are in the same scale.

##### Violin plots

We used violin plot function (https://github.com/bastibe/Violinplot-Matlab) to make all our violin plots.

##### Alluvial flow diagram

We used alluvialflow function to generate the diagram in Figure S5C (https://www.mathworks.com/matlabcentral/fileexchange/66746-alluvial-flow-diagram).

### Statistics

A power analysis was not performed a-priori to determine sample size. All statistical analyses were performed in Matlab, using non-parametric tests with significance levels set at α < 0.05, and did Bonferroni corrections on α for multiple comparisons when necessary. Mann-Whitney U-tests (Wilcoxon rank sum test) were used to test differences between two independent populations. When comparing more than two populations that were non-normally distributed, a Kruskal-Wallis test, a nonparametric version of one-way ANOVA, was used to determine whether statistically significant differences existed among these independent populations. If significant differences did exist, post hoc multiple comparison tests (multcompare in Matlab) with Bonferroni correction were used to test for significant differences between pairs within the group. To compare more than two proportions (Figure 1F), Tukey’s HSD multiple comparisons test among proportions were used (https://www.mathworks.com/matlabcentral/fileexchange/15499-tmcomptest). For pairwise comparisons of proportions (Figure 6B), Chi-square test was used (https://www.mathworks.com/matlabcentral/fileexchange/45966-compare-two-proportions-chi-square).

### Code and data availability

All analyses and plotting were performed in Matlab. The databases for each set of experiment and corresponding analysis and plotting codes are available upon request.

---

**Figure S1.**
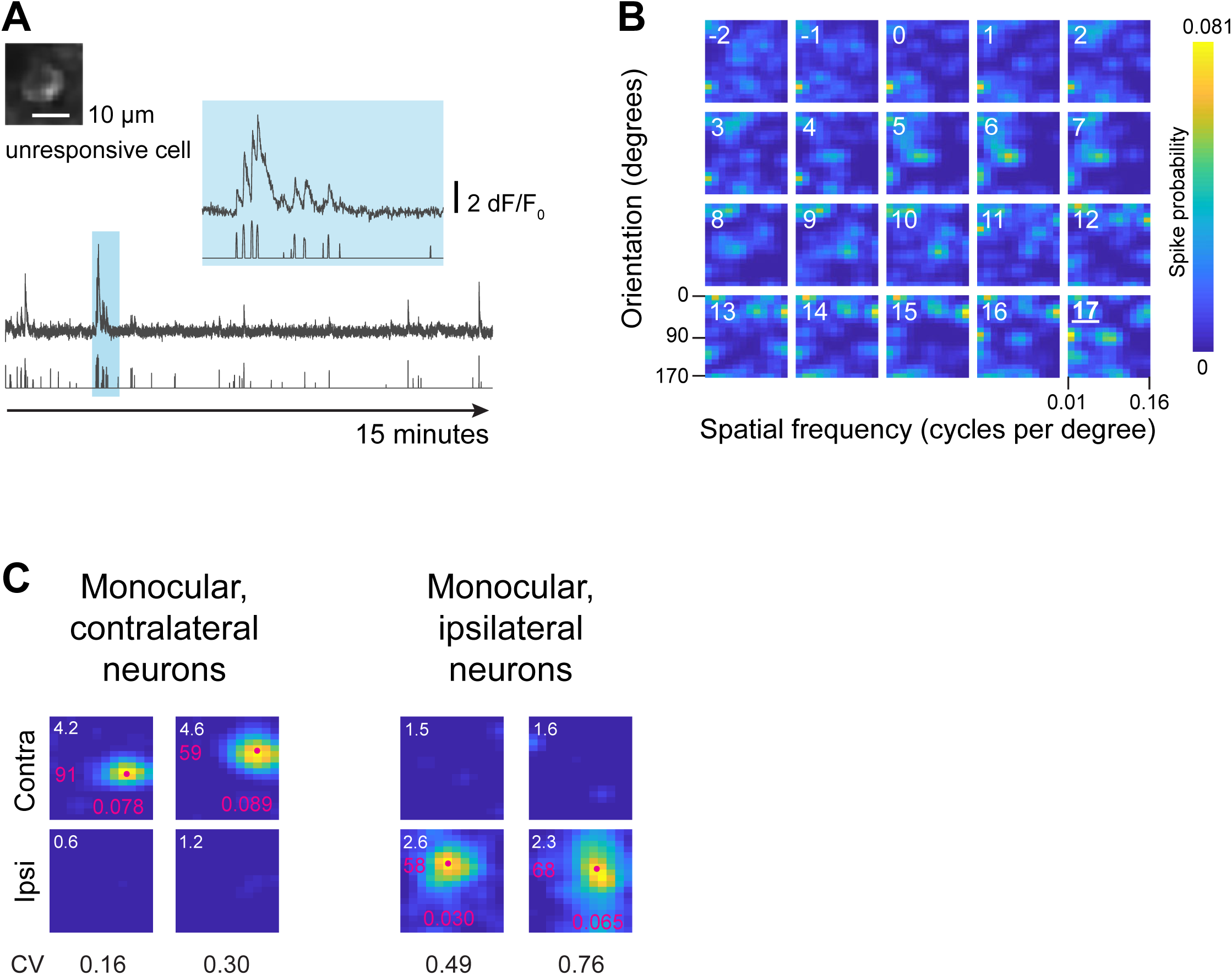
Example of unresponsive and monocular neurons, plus examples of tuning measurements. Related to Figure 1. **(A)** Fluorescence changes and estimated spike rate for a visually unresponsive neuron. The cell is shown in the upper left. The blue highlighted region has been expanded horizontally and shown above for clarity. Scale bar for fluorescence changes is shown to the right of this expanded trace. Example is from responses to contralateral eye stimulation. **(B)** Spike probability as a function of stimulus onset for the cell in panel A. Maps are derived from 18 orientations at 12 spatial frequencies presented 16 to 17 times. For this cell, optimal delay occurs 17 imaging frames, or 1.097 second after stimulus onset. **(C)** Example tuning kernels for two monocular contralateral and monocular ipsilateral neurons, respectively. Tuning kernels to contralateral eye are in upper panel, and ipsilateral in lower panel. SNR is shown in the upper left corner of each kernel. Orientation and spatial frequency tuning preference is marked with a magenta dot and written along each axis (orientation on the y-axis, spatial frequency on the x-axis). Circular variance values (CV) for each neuron are shown at the bottom.

**Figure S2.**
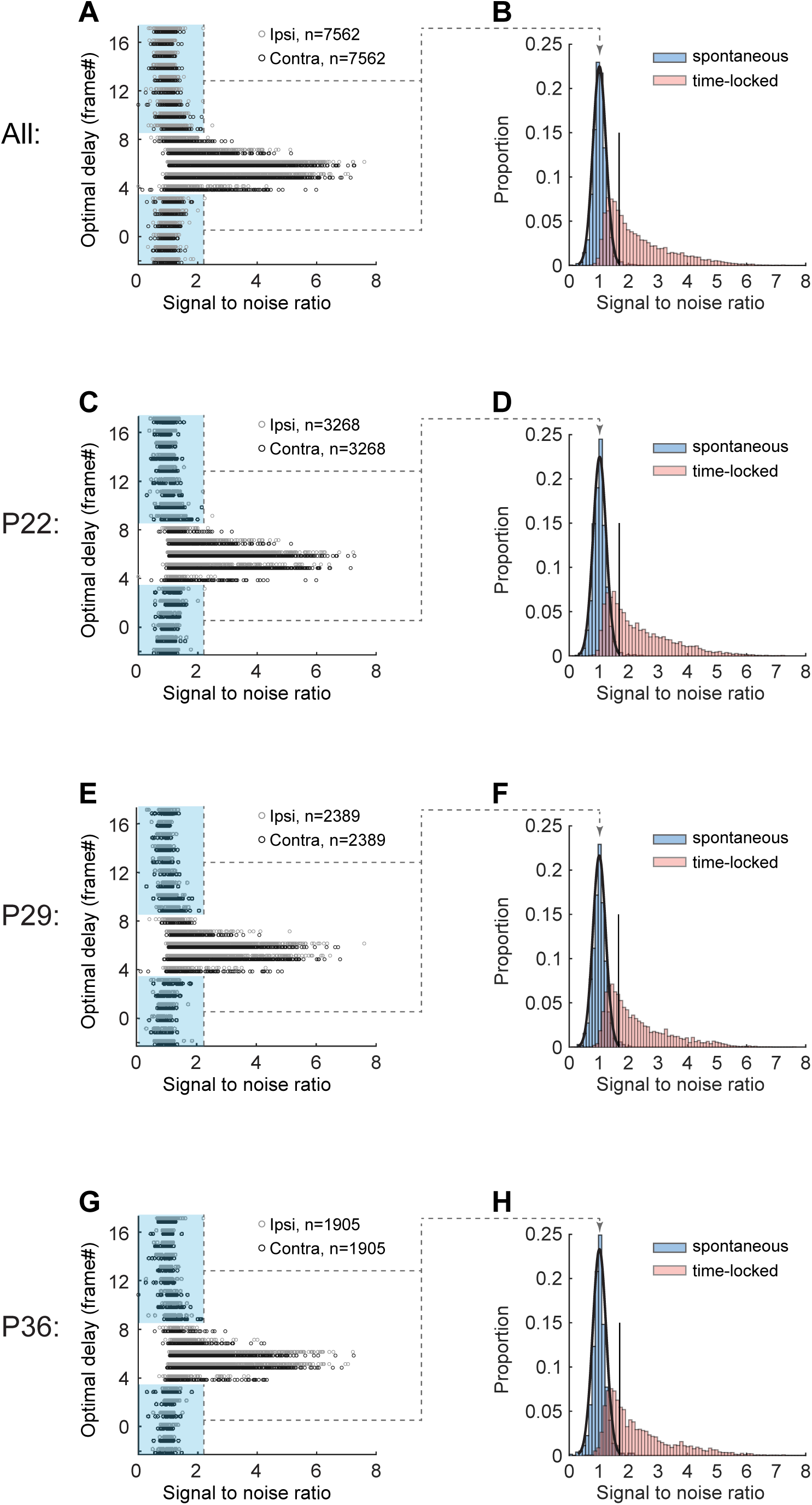
SNR distribution of all neurons imaged at three times in NR mice. Related to Figure 1. **(A)** Plot of SNR as a function of optimal delay for all 7562 L2/3 neurons imaged across the critical period in NR mice. Dark and light gray represent responses to contralateral and ipsilateral eye stimulation, respectively. Cells whose optimal delay were -2 to 3 and 9 to 17 frames post stimulus onset are visually unresponsive (blue shading). **(B)** Blue shaded histogram and normal distribution fit of SNR values for unresponsive neurons highlighted by the blue rectangles in panel A. Black vertical line denotes the threshold of responsiveness: 3 standard deviations above the mean of this normal distribution. Red shaded histogram: neurons whose spiking occurred between 4 and 8 frames post stimulus. Only neurons with optimal delays in this window that also had an SNR value above the threshold were considered responsive. This would be the subset of the red-shaded histogram that is to the right of the vertical line. (**C and D**) Same plots as in A and B, but for L2/3 neurons imaged at P22 in NR mice. (**E and F**) Same plots as in A and B, but for L2/3 neurons imaged at P29 in NR mice. (**G and H**) Same plots as in A and B, but for L2/3 neurons imaged at P36 in NR mice. Note that the distribution of SNR and the position of the vertical line across three times are highly similar.

**Figure S3.**
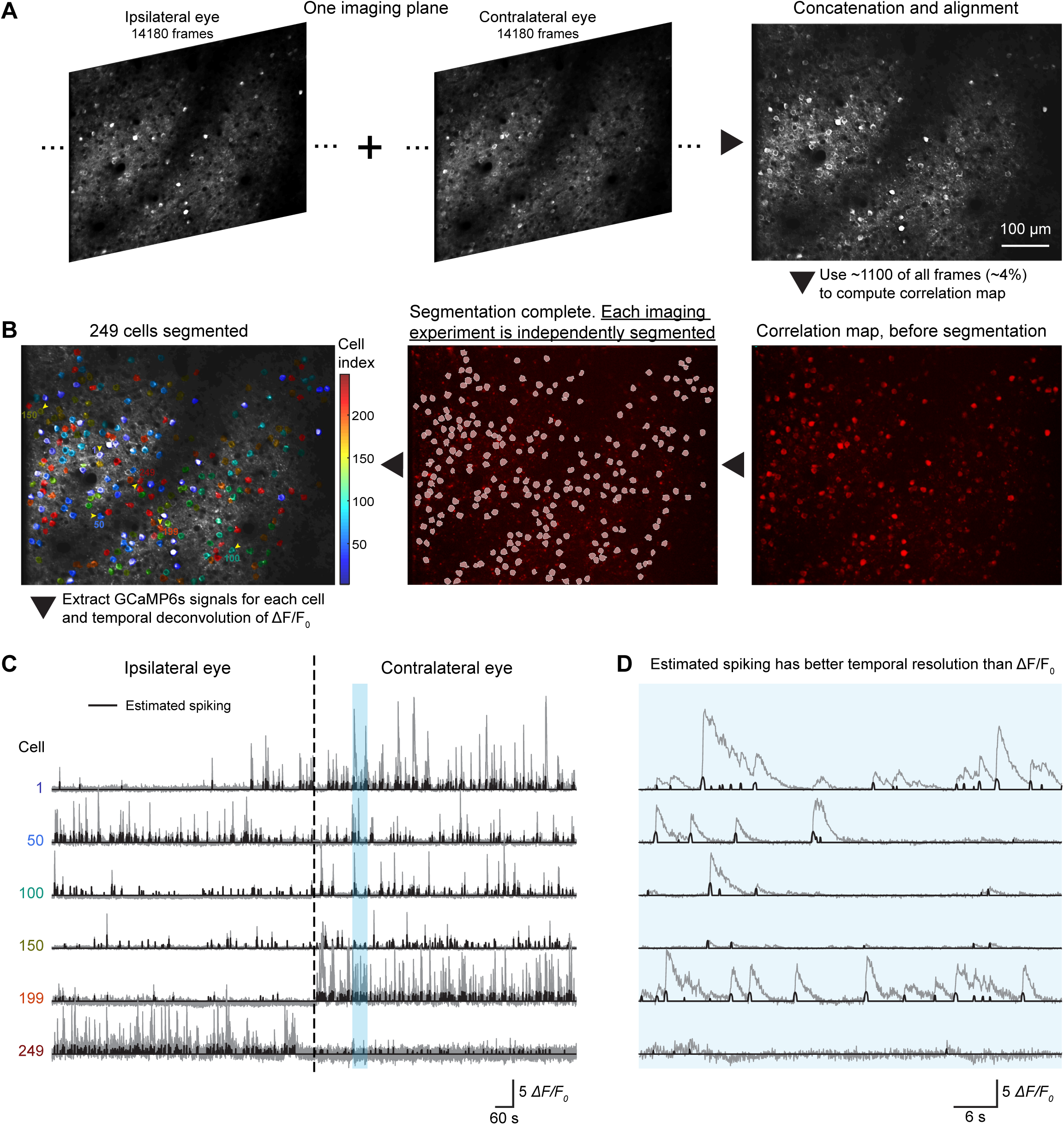
Example of cell segmentation, signal extraction and deconvolution for a single plane imaged from visual stimulation to two eyes separately. Related to Figure 2. **(A)** Time series data of responses recorded via stimulation of the ipsilateral and contralateral eye. Data sets were concatenated into one file and then motion corrected (concatenation and alignment). **(B) Right**: Correlation map calculated from ∼1,100 frames of images in the concatenated and aligned image series file. Note only ∼4% of total image frames were used for segmentation. This was limited by GPU performance (Nvidia Quadro M5000). **Center**: 249 cells segmented from the correlation map. **Left**: Overlay of motion-corrected average fluorescence image and segmentation masks from the middle panel. To ease visualization, segmented cells are colored by their indices, which are given from the order of their segmentation. The responses plotted in panels C and D are from neurons identified by the yellow arrowheads in this panel. (**C and D**) Examples of fluorescence changes (ΔF/F0) and estimated spike rates over consecutive imaging sessions of ipsilateral and contralateral eye stimulation. Responses for the 6 cells marked by yellow arrowheads in the left panel of B are shown. For each cell, the ΔF/F0 series is plotted in gray and the estimated spike rate in black. The time highlighted by the light blue bar is expanded horizontally in D.

**Figure S4.**
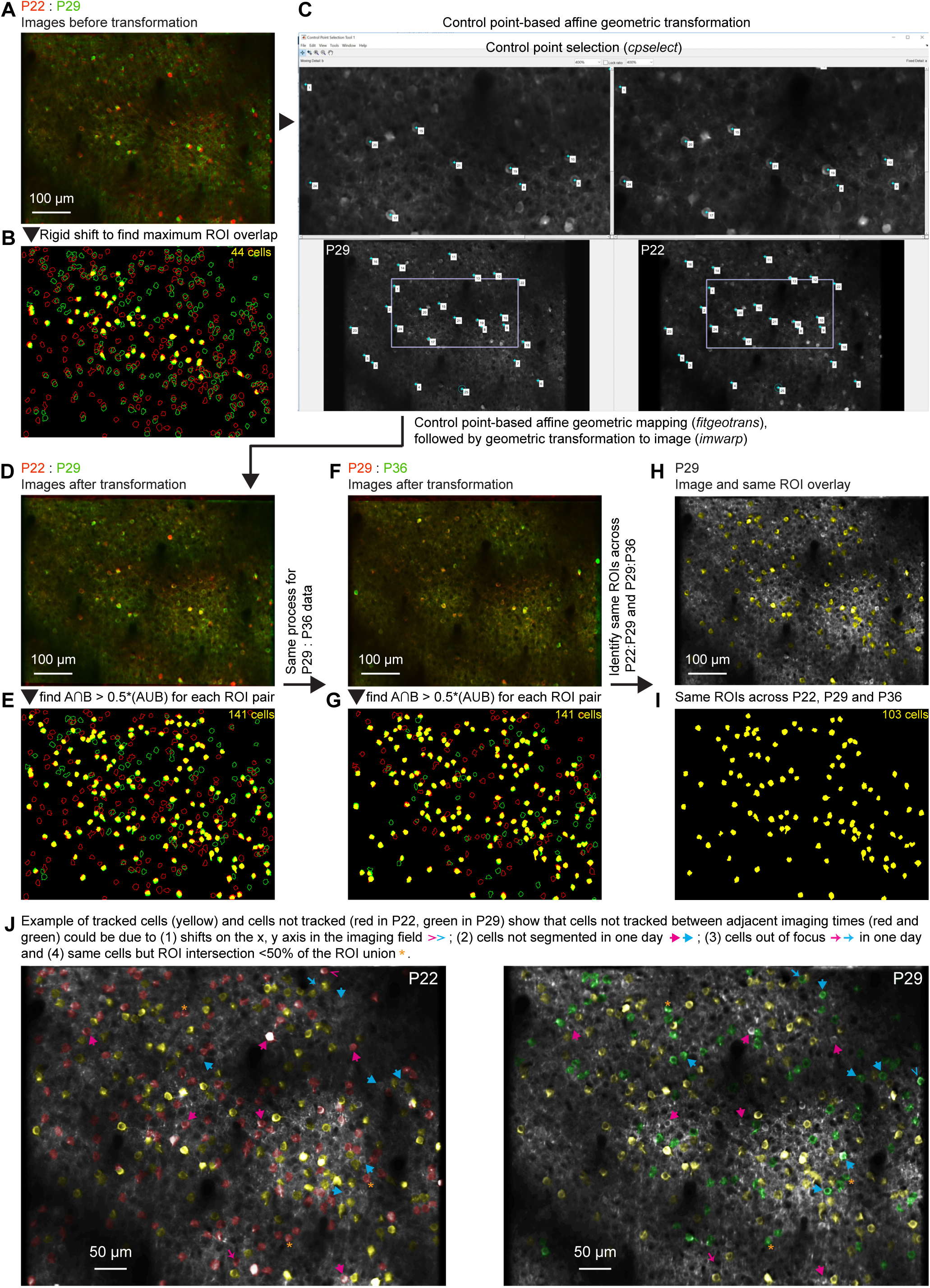
Example of tracked cell identification across days for one imaging plane and explanation of cells failed to be tracked between days. Related to Figure 2. **(A)** Overlay of average fluorescence images of a focal plane longitudinally imaged on P22 (red) and P29 (green) before image transformation. **(B)** Identification of 44 cells (yellow shaded ROIs) tracked between P22 and P29 before image transformation. The criterion for tracked cell identification is ROI intersection over 50% of ROI union. ROI outlines are cell segmentations following protocol shown in Figure S3. Red and green outlines represent cells failed to be tracked, with red marking segmentations from P22 imaging, green marking segmentations from P29 imaging. **(C)** Example of doing control point selection on two images shown in panel A for subsequent affine geometric transformation. Note the similarity in the pattern of cells at P22 and P29. **(D)** Overlay of images in panel A after control point-based affine geometric image transformation. Note the improvement from A. **(E)** Identification of 141 cells (yellow shaded ROIs) tracked between P22 and P29 after geometric image transformation, with the same criterion for tracked cell identification as in B. Note the marked increase on number of tracked cells compared to B (before image transformation). Red and green outlines represent cells failed to be tracked, with red marking segmentations from P22 imaging, green marking segmentations from P29 imaging. **(F)** Overlay of average fluorescence images of the plane in panel D longitudinally imaged at P29 (red) and P36 (green), after control point-based affine geometric image transformation. **(G)** Identification of 141 cells (yellow shaded ROIs) tracked between P29 and P36 after image transformation. Red and green outlines represent cells failed to be tracked, with red marking segmentations from P29 imaging, green marking segmentations from P36 imaging. (**H and I**) Identification of 103 cells tracked across P22-P36, by finding intersection of cells between E and G. Panel H shows the overlay of fluorescence image at P29 and ROIs of tracked cells in yellow. Panel I shows just the ROIs of the tracked cells in yellow. (**J**) Explanation of cells failed to be tracked between days using P22 and P29 as an example. Cells tracked between P22 and P29 are colored with yellow masks in the two fluorescence images. Cells segmented in P22 and failed to be tracked are colored with red masks in P22 image; cells segmented in P29 and failed to be tracked are colored with green masks in P29 image. Cells without color masks (white) in each image are cells not segmented. Comparing the two images revealed that cells failed to be tracked between adjacent imaging times (red and green labeled cells) could be due to: (1) shifts on the x, y-axis in the imaging field (> marker close to image edge), (2) cells not segmented in one day but segmented in the other day (thick arrow marker), (3) cells out of focus on a given session (thin arrow marker), (4) same cells but ROI intersection <50% of the ROI union (* marker). For reasons 2 and 3, cells segmented in P22 but not P29 are identified by magenta markers in both images, cells segmented in P29 but not P22 are identified by cyan markers in both images. For reason 4, cells are marked by * with orange color in both images.

**Figure S5.**
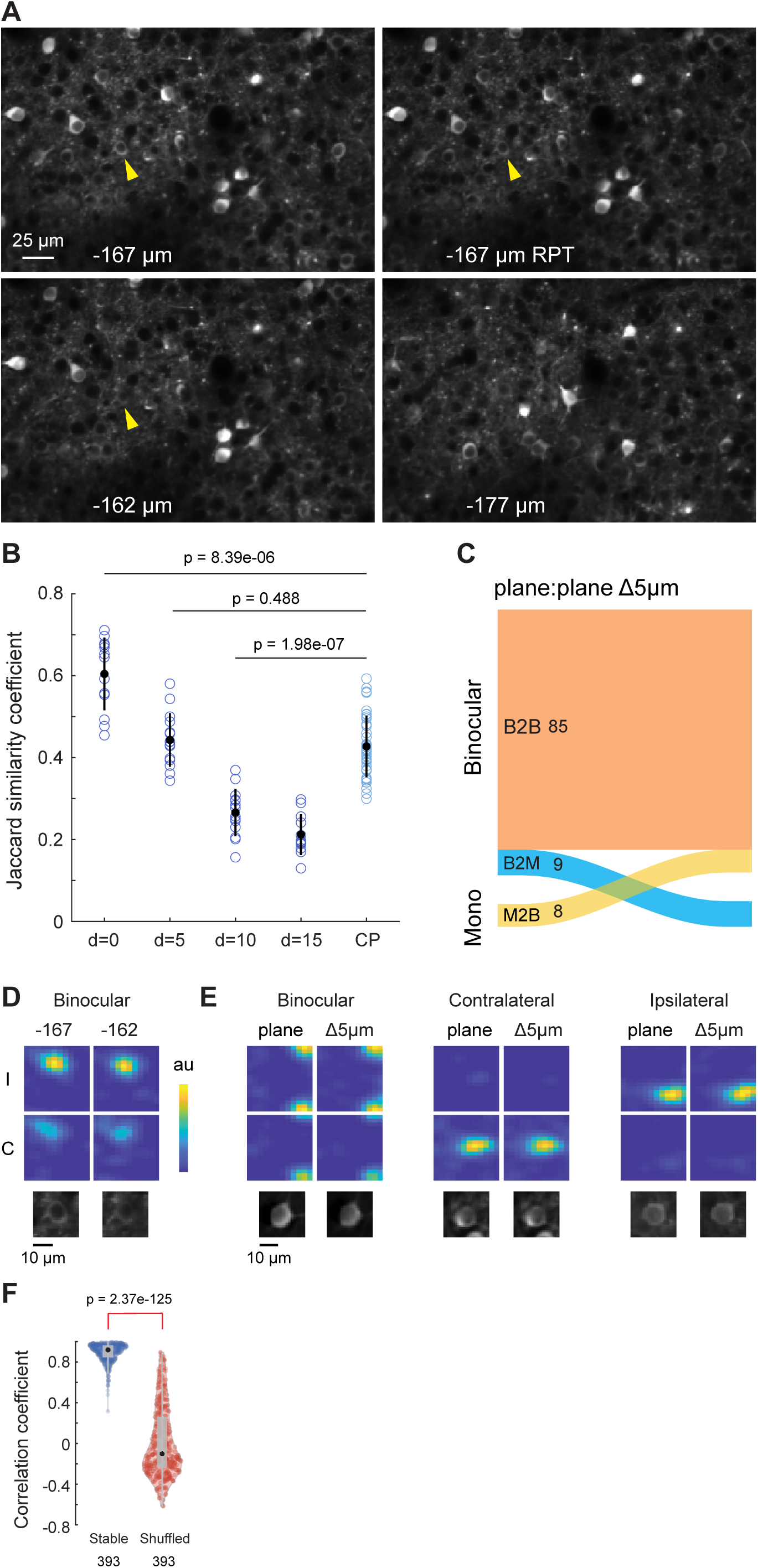
Effect of focal plane drift on tuning measures and binocularity. Related to Figure 2. **(A)** Partial field of view of four consecutively imaged planes at various depths: -167μm below pia (−167μm), the same depth (−167μm RPT), 5μm shallower (−162μm) and 10μm deeper (−177μm). Note the similarity of the top two images (Δ0μm in focal plane depth), and their similarity to the bottom left image (Δ5μm in focal plane depth). Note also the difference from the bottom right (Δ10μm in focal plane depth). Yellow arrowheads in the first three images point to the same cell. Its tuning kernels, measured at 0 and 5 um focal displacement, are shown in D. **(B)** Plot of Jaccard similarity coefficients as a function of focal plane displacement (d=0 to d=15); also plotted are these coefficients derived from the layer 2/3 neurons in Figure 3-5 that were longitudinally imaged across the critical period (CP) in normally and dark-reared mice. Black dot and line indicate the mean and standard deviation. Number of data points from left to right are: 13, 15, 14, 12, and 40. Mann-Whitney U test was used for comparing between two groups of data, p-values are indicated for comparisons (α=0.005 after Bonferroni correction for pairwise comparison). Note that the distribution at CP is the same as d=5, indicating that focal plane differences of longitudinal imaging is no bigger than 5μm. **(C)** Alluvial flow diagram for binocular neurons in experiments with 5μm focal plane displacement. Thickness of the flows indicates the number of cells. Stable binocular (B2B) = 85, binocular to monocular (B2M) = 9, monocular to binocular (M2B) = 8. Note the turnover of binocular neuron is less than 10% (9/ (85+9)), which is markedly lower than what we find in either our critical period or adult longitudinal data sets (see Figure 6B). **(D)** Soma and tuning kernels of a stable binocular cell marked in A. Top and bottom rows show responses to ipsilateral and contralateral eyes, respectively. **(E)** Examples of three more stable binocular and monocular cells imaged at 5μm focal plane displacement. For D and E, Color indicates estimated spiking at each possible combination of orientation and spatial frequency. Color map of estimated spiking is normalized for each cell. **(F)** Violin plots of correlation coefficients of receptive field tuning kernels for the same cells imaged with 5μm focal plane displacement (stable) and for random pairings of cells (shuffled). The tapered tail extending towards higher coefficients in the shuffled control reflects random pairing of kernels with high circular variance (>0.7) and low spatial frequency preference (<0.02 cycles/degree). P-value is derived from Mann-Whitney U test. Number of samples in each group is shown below.

**Figure S6.**
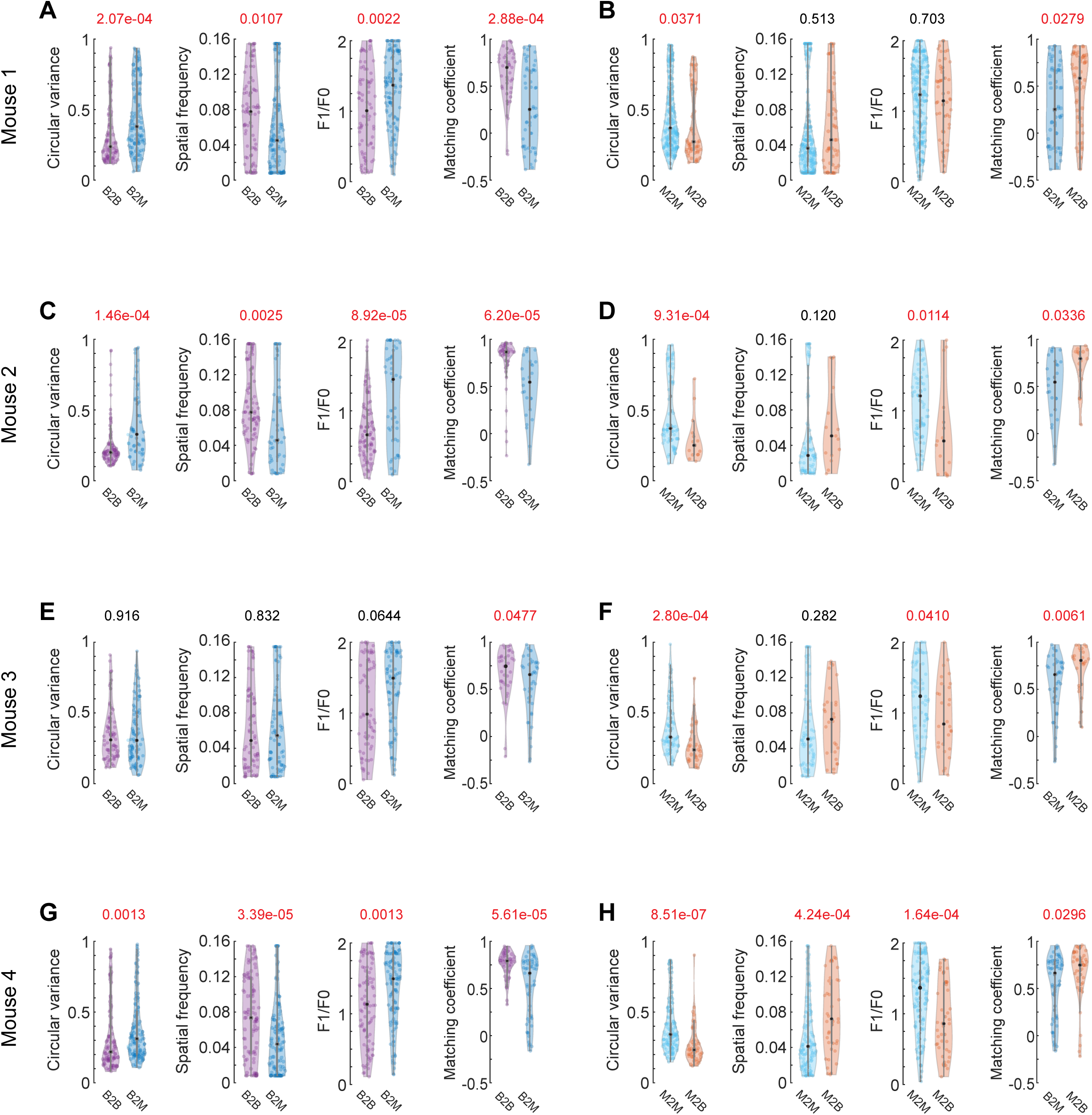
Quantification of layer 2/3 receptive field tuning properties in four normally reared mice during the critical period. Related to Figure 3 and 4. **(A)** Violin plots of receptive field tuning measurements for binocular neurons in mouse 1. Plotting and statistics are as in Figure 3C. B2B, n =41; B2M, n = 56. **(B)** Violin plots of receptive field tuning measurements for monocular contralateral neurons in mouse 1. Plotting and statistics as in Figure 4C. M2B, n = 52; M2M, n = 234. **(C)** Same plots as A, but for mouse 2. B2B, n =44; B2M, n = 26. **(D)** Same plots as B, but for mouse 2. M2B, n = 15; M2M, n = 76. **(E)** Same plots as A, but for mouse 3. B2B, n =35; B2M, n = 41. **(F)** Same plots as B, but for mouse 3. M2B, n = 28; M2M, n = 74. **(G)** Same plots as A, but for mouse 4. B2B, n =50; B2M, n = 59. **(H)** Same plots as B, but for mouse 4. M2B, n = 43; M2M, n = 132.

**Figure S7.**
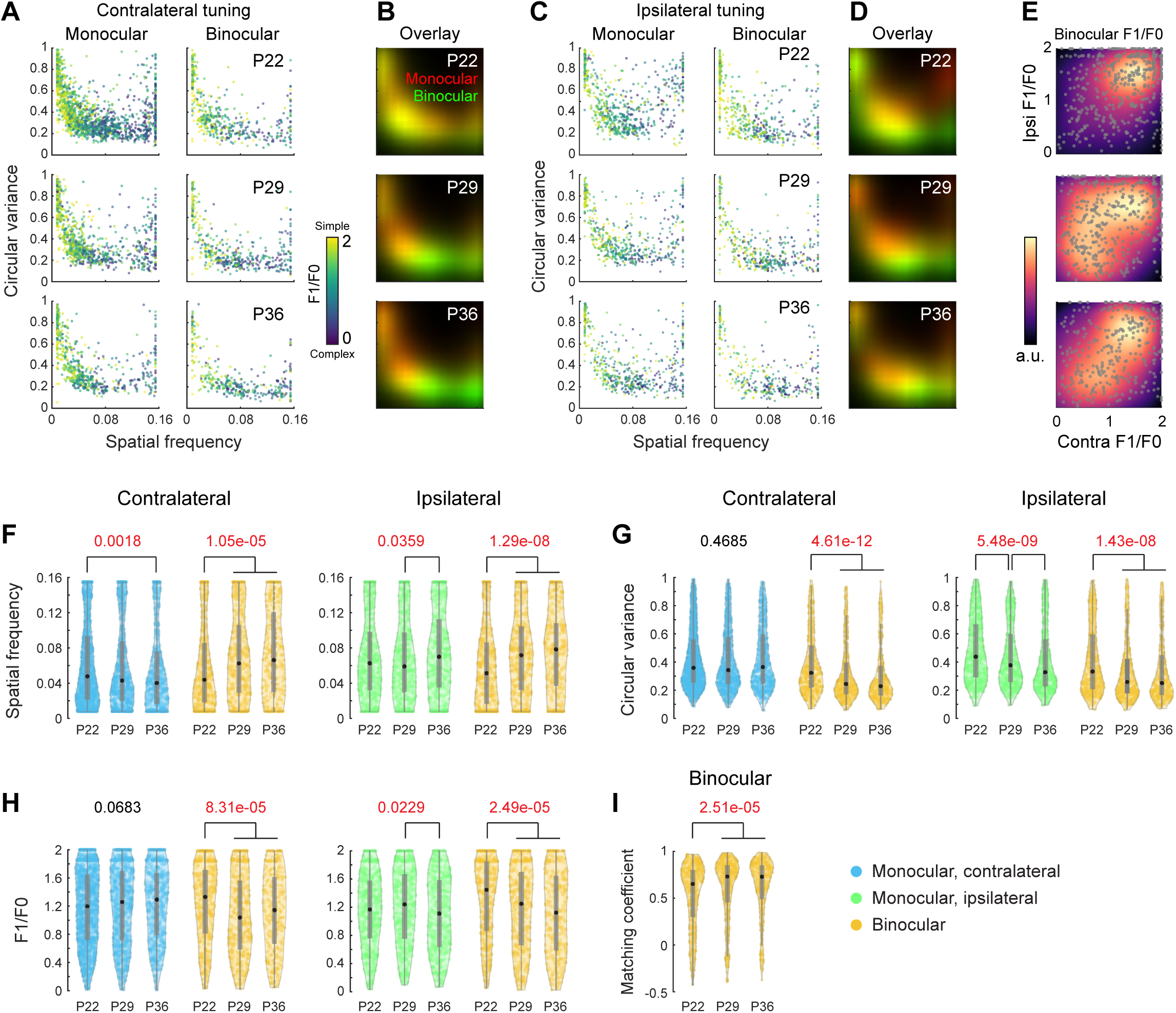
Improvement of ipsilateral and binocular receptive field tuning in layer 2/3 in normally reared mice during the critical period. Related to Figure 3 and 4. **(A)** Plots of circular variance (ordinate), spatial frequency preference (abscissa) and phase invariance, measured as F1/F0 (color bar) for all layer 2/3 cells imaged at P22 (top row), P29 (middle row), and P36 (bottom row). Left columns plots these values for cells that were monocular contralateral at these ages. Right column plots these values for contralateral eye responses of binocular neurons. P22: 4 mice, 1444 monocular contra, 451 binocular; P29: 4 mice, 833 monocular contra, 418 binocular; P36, 4 mice, 584 monocular contra, 339 binocular. **(B)** Density profiles of monocular and binocular responses in panel A overlaid in red and green. Note the progressive separation of the binocular from monocular pools with binocular neurons (green) becoming increasingly tuned to higher spatial frequencies (rightward shift) and lower circular variance (downward shift). **(C)** Same plots as in A, but for responses to ipsilateral eye. P22: 574 monocular ipsi, 451 binocular; P29: 515 monocular ipsi, 418 binocular; P36, 495 monocular ipsi, 339 binocular. **(D)** Density profiles of monocular and binocular responses in panel C overlaid in red and green. **(E)** F1/F0 plots of binocular neurons. Ordinate plots F1/F0 for ipsilateral eye evoked responses of binocular neurons. Abscissa plots this for contralateral eye evoked responses. Color bar is in arbitrary units and represents density. Upper right quadrant is populated by “simple” cells. Lower left quadrant by “complex” cells. Note that binocular neurons at P22 are largely simple, but by P36 the full distribution of complex and simple receptive fields are present in the binocular pool. **(F)** Violin plots of spatial frequency preference in panels A and C. Monocular responses are plotted adjacent to same-eye responses recorded from binocular neurons. Black dot, median; gray vertical lines, quartiles with whiskers extending to 2.698σ. Statistics: Kruskal-Wallis one-way analysis of variance, with p-values shown above each group and significant values in red color. Black brackets denote significance using multiple comparison test with Bonferroni correction for pairwise comparisons. **(G)** As in F, but for circular variance. **(H)** As in F, but for F1/F0. **(I)** Violin plots of binocular matching coefficient for all L2/3 binocular neurons at each age. Statistics: Kruskal-Wallis one-way analysis of variance, followed by multiple comparison test with Bonferroni correction.

**Figure S8.**
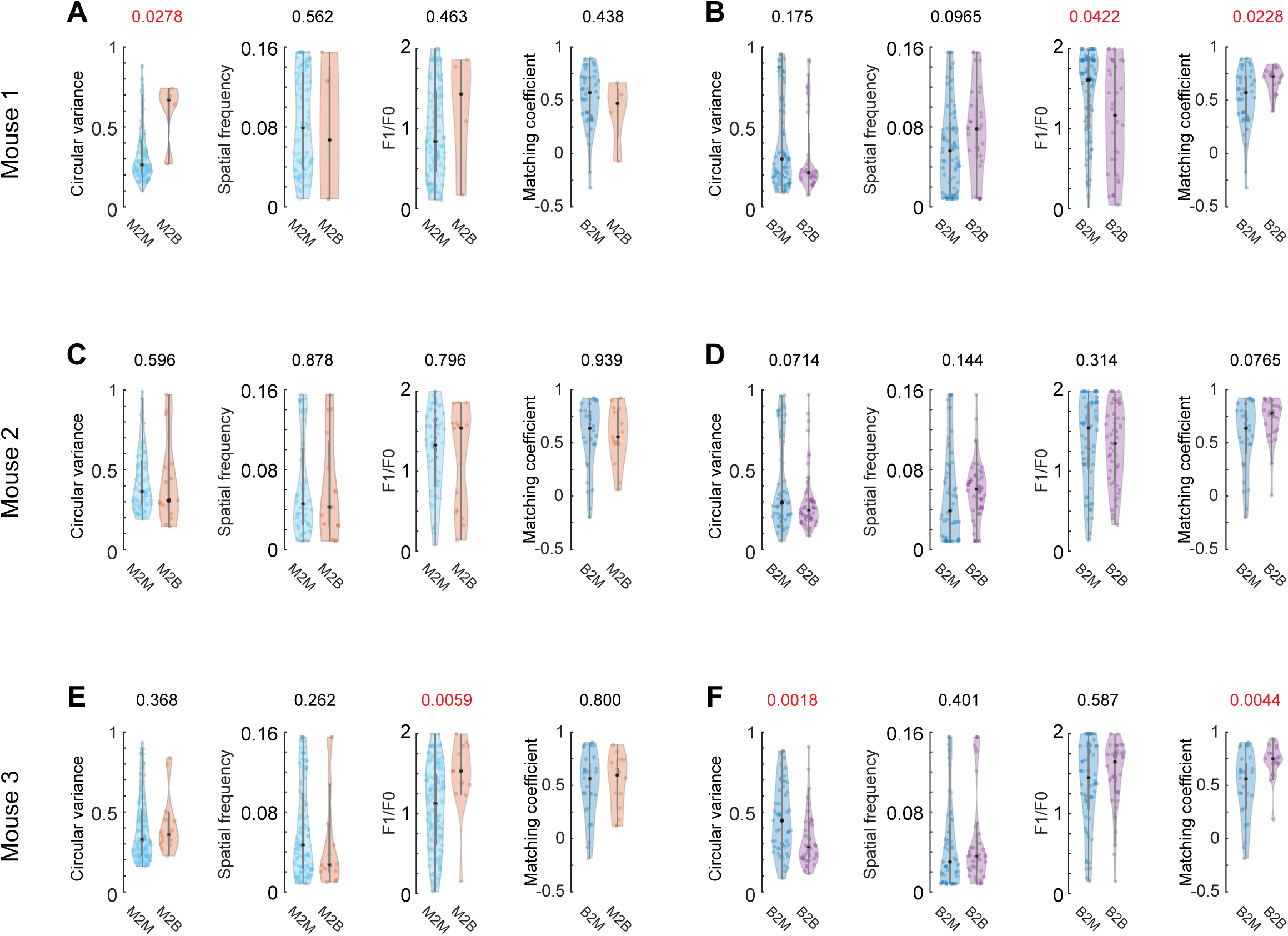
Quantification for tuning properties in three individual mice dark-reared during the critical period. Related to Figure 5. **(A)** Violin plots of receptive field tuning measurements for monocular contralateral neurons in dark-reared (DR) mouse 1. Plotting and statistics are done the same as in Figure 5A. M2B, n = 4; M2M, n = 105. **(B)** Violin plots of receptive field tuning measurements for binocular neurons in DR mouse 1. Plotting and statistics are done the same as in Figure 5B. B2B, n =16; B2M, n = 40. **(C)** As in A, but for DR mouse 2. M2B, n = 15; M2M, n = 65. **(D)** As in B, but for DR mouse 2. B2B, n =24; B2M, n = 28. **(E)** As in A, but for DR mouse 3. M2B, n=17; M2M, n = 135. **(F)** As in B, but for DR mouse 3. B2B, n = 19; B2M, n=28.

**Figure S9.**
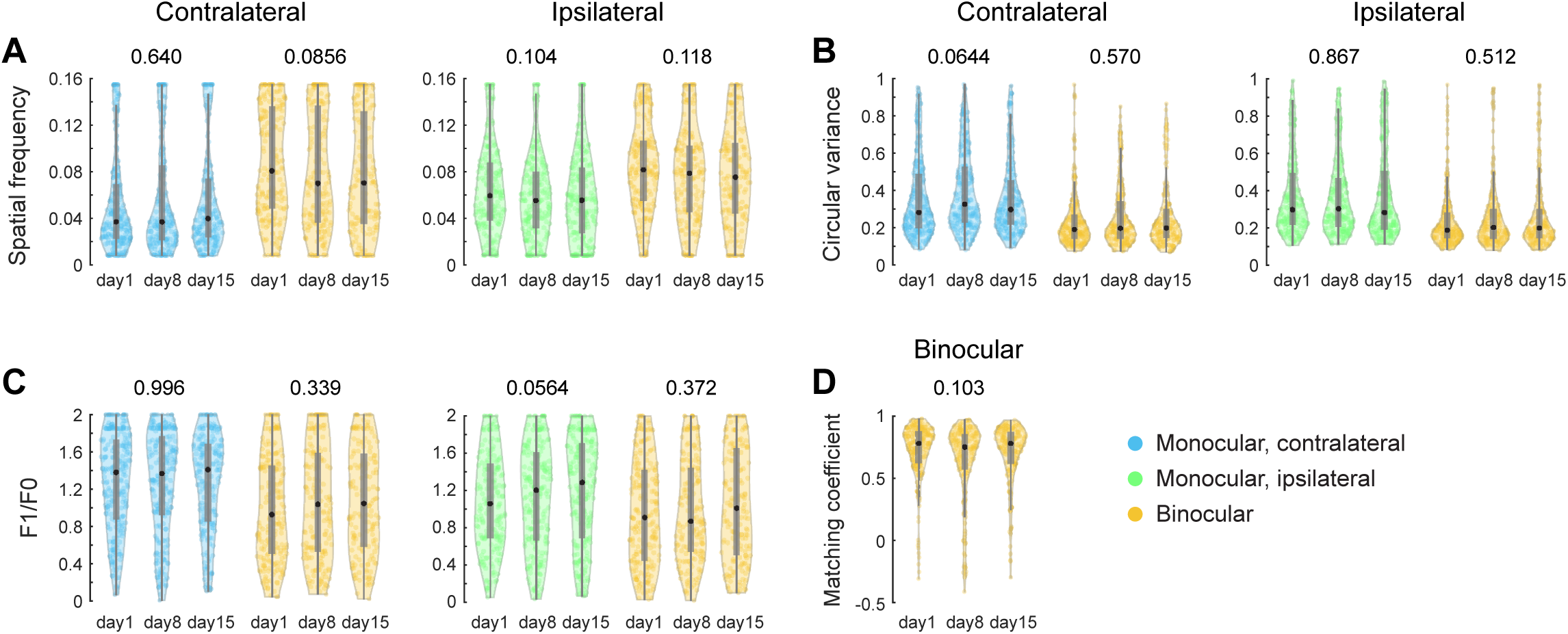
No change of receptive field tuning in layer 2/3 in adult mice. Related to Figure 6. **(A)** Violin plots of spatial frequency preference for all cells imaged over 2-week period in three adult mice. Plots and statistical analysis as in Figure S7F. Day1: 254 monocular contra, 197 monocular ipsi, 210 binocular; Day8: 231 monocular contra, 213 monocular ipsi, 182 binocular; Day15, 204 monocular contra, 225 monocular ipsi, 167 binocular. **(B)** As in A, but for circular variance. **(C)** As in A, but for F1/F0. **(D)** Violin plots of binocular matching coefficient for all adult L2/3 binocular neurons at each imaging day. Statistics: Kruskal-Wallis one-way analysis of variance.

**Figure S10.**
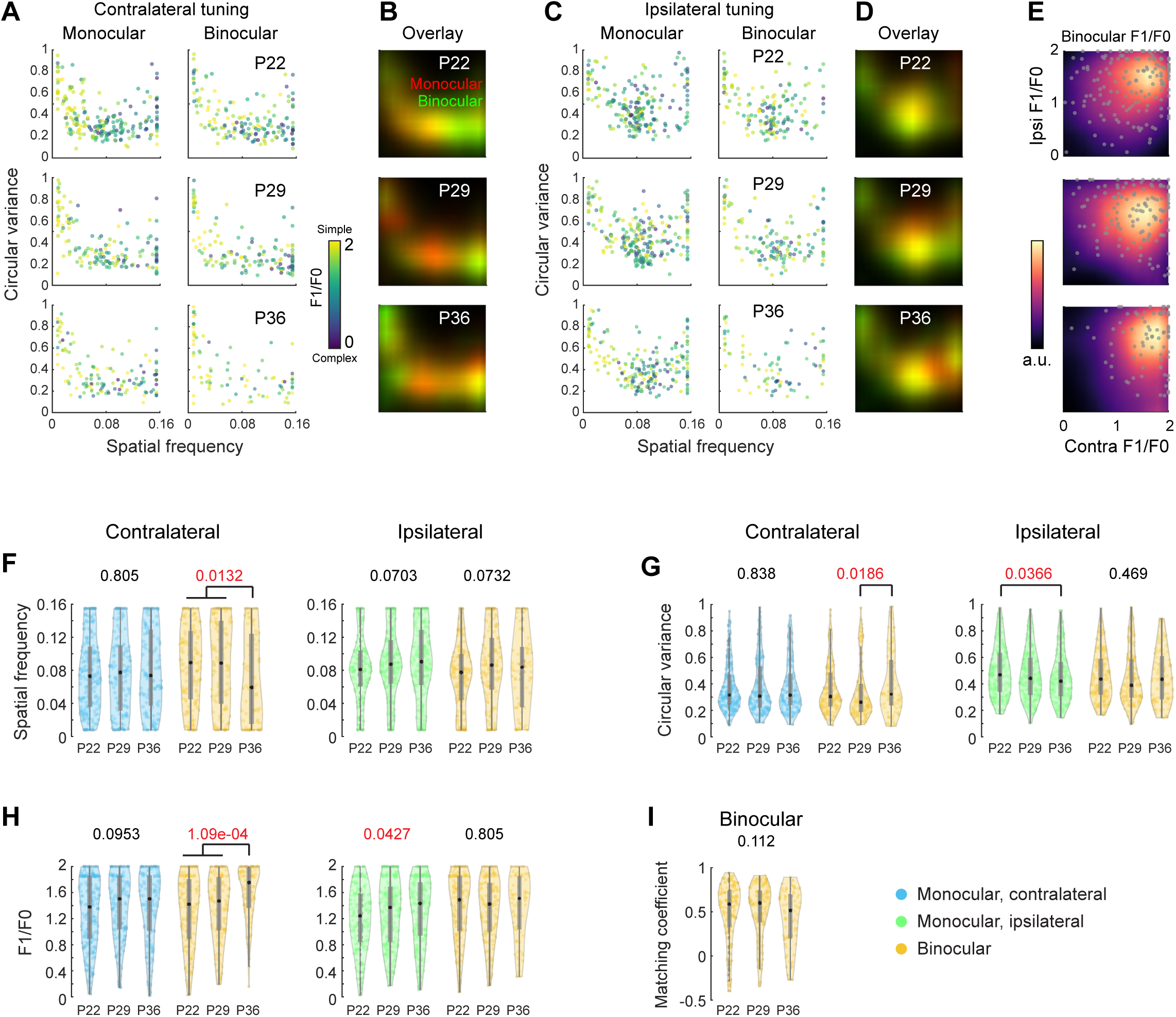
No improvement of receptive field tuning in layer 4 during the critical period. Related to Figure 7. **(A)** Plots of circular variance (ordinate), spatial frequency preference (abscissa) and phase invariance, measured as F1/F0 (color bar) for 717 longitudinally tracked layer 4 cells from 4 mice at P22 (top row), P29 (middle row), and P36 (bottom row). Plots are as in Figure S7A. P22: 212 monocular contra, 155 binocular; P29: 146 monocular contra, 132 binocular; P36, 130 monocular contra, 85 binocular. **(B)** Density profiles of monocular and binocular responses in panel A overlaid in red and green. **(C)** Plots as in A, but for responses to the ipsilateral eye. P22: 195 monocular ipsilateral, 155 binocular; P29: 220 monocular ipsilateral, 132 binocular; P36, 205 monocular ipsilateral, 85 binocular. **(D)** Density profiles of monocular and binocular responses in panel C overlaid in red and green. **(E)** F1/F0 plots of longitudinally tracked layer 4 binocular neurons, plotted as in Figure S7E. Upper right quadrant is populated by “simple” cells. Lower left quadrant by “complex” cells. Note that layer 4 binocular neurons are largely simple and a regression of the density pattern for the contra F1/F0. **(F)** Violin plots of spatial frequency preference in panels A and C. Monocular responses are plotted adjacent to same-eye responses recorded from binocular neurons. Black dot, median; gray vertical lines; quartiles with whiskers extending to 2.698σ. Statistics: Kruskal-Wallis one-way analysis of variance, with p-values shown above each group and significant values in red color. Black brackets denote significance using multiple comparison test with Bonferroni correction for pairwise comparisons. **(G)** As in F, but for circular variance. **(H)** As in F, but for F1/F0. **(I)** Violin plots of binocular matching coefficient for layer 4 binocular neurons among longitudinally imaged cells at each age. Statistics: Kruskal-Wallis one-way analysis of variance, followed by multiple comparison test with Bonferroni correction.

